# Therapeutic targeting of fibrin–microglia interactions ameliorates Alzheimer’s disease-related hyperexcitability and brain network dysfunction

**DOI:** 10.64898/2026.05.01.722324

**Authors:** Kelli Lauderdale, Zhaoqi Yan, Andrew S. Mendiola, Yutong Zhang, Dakota Mallen, Pranav Nambiar, Erica Brady, Stephanie R. Miller, Rosa Meza Acevedo, Belinda Cabriga, Fred Jiang, Nick Kaliss, Kevin Shen, Jia Shin, Jessica Herbert, Keran Ma, Jae Kyu Ryu, Ayushi Agrawal, Renaud Schuck, Maria del Pilar S. Alzamora, Jorge Sanz-Ros, Inma Cobos, Jeffrey Stavenhagen, Aaron B. Kantor, Mark H. Ellisman, Katerina Akassoglou, Jorge J. Palop

**Affiliations:** Gladstone Institute of Neurological Disease, San Francisco, CA 94158, USA; Department of Neurology, Weill Institute for Neurosciences, University of California, San Francisco, CA 94158, USA; Center for Neurovascular Brain Immunology at Gladstone and UCSF, San Francisco, CA 94158; Department of Pharmacology, University of California, San Diego, La Jolla, CA 92093, USA; Neuroscience Graduate Program, UCSF, San Francisco, California, USA; Department of Neurobiology and Anatomy, McGovern Medical School, University of Texas Health Science Center at Houston, Houston, TX, 77030; Gladstone Institute of Data Science and Biotechnology; San Francisco, CA 94158, USA; Department of Pathology, Stanford University School of Medicine, Stanford, CA 94305, USA; Therini Bio, 2108 N ST, Ste 4963, Sacramento, CA 95816, USA; Department of Neurosciences, School of Medicine, University of California, San Diego, La Jolla, CA 92093, USA

**Author notes:** Equal co-second author contribution (Zhaoqi Yan, Andrew S. Mendiola, Yutong Zhang, Dakota Mallen).

## Abstract

Brain network dysfunction—including hyperexcitability, altered oscillations, and sleep disruption—is prominent in Alzheimer’s disease (AD), but the contribution of vascular-neuroimmune processes to these alterations remains unclear. Here, we blocked the pro-inflammatory interaction of the blood protein fibrin with microglia using genetic (Fgg^γ390–396A^ mice) and antibody-based (5B8 and THN392) strategies to test its role in AD-related network dysfunction. The 5xFAD model of AD exhibited network hyperexcitability associated with oscillatory slowing, sleep states, and disrupted sleep-circadian rhythms. These deficits were largely attenuated by blocking fibrin-microglia interactions in 5xFAD;Fgg^γ390–396A^ mice. Notably, pharmacological interventions after disease onset with both anti-fibrin antibodies similarly attenuated these AD-related network deficits and behavioral abnormalities. We conclude that vascular-neuroimmune processes driven by fibrin-microglia interactions promote AD-related network dysfunction and that targeting the fibrin-microglia axis—currently under clinical evaluation with the humanized antibody THN391— represents a promising therapeutic strategy for AD. There is a companion manuscript submitted to *bioRxiv* (Yan et al., 2026).^110^

**HIGHLIGHTS:**

- Fibrin promotes AD-related network hyperexcitability and oscillatory slowing in 5xFAD mice
- Fibrin promotes AD-related disruption of sleep and circadian rhythms in 5xFAD mice
- Genetic blocking of fibrin-microglia interactions rescues AD-related brain network dysfunction
- Anti-fibrin antibodies (5B8 and THN392) show acute and chronic therapeutic benefit

## INTRODUCTION

Growing evidence indicates that vascular dysfunction is central to AD pathogenesis.^1, 2^ A diverse array of cerebrovascular abnormalities—including blood-brain barrier disruption (BBBd), impaired hemodynamics, and neurovascular unit dysregulation—underscores the critical role of vascular health in AD progression.^3–6^ These pathological changes significantly impact neurovascular coupling, brain homeostasis, metabolism, and the clearance of toxic metabolites and Aβ.^7–9^ Increased fibrinogen in the CSF and plasma represents an early marker across the general AD population that correlates with AD biomarkers of disease progression, including Aβ, tau, and cognitive decline.^10–13^ Fibrinogen levels also increase progressively with normal aging in humans and mice,^14, 15^ suggesting that age-related thromboinflammatory changes may also play a role in cognitive aging. Notably, a recent cerebrospinal fluid (CSF) proteomics study has identified five molecular subtypes of AD pathogenesis, including a BBBd-dominant subtype.^16^ In this subtype, fibrinogen and other blood proteins are elevated in the CSF, mapping onto pathways involving coagulation, immunity, and inflammatory response. The BBBd subtype also exhibits reduced CSF levels of neuroplasticity-related proteins converging on the transcription factor REST, which is implicated in aging and neurodegeneration.^17^ These findings suggest that the BBB is compromised and that blood protein leakage triggers blood-driven inflammatory and synaptic responses. Altogether, these clinical findings suggest that fibrin(ogen) and vascular dysfunction define a distinct pathological and molecular signature that explains a significant proportion of the heterogeneity within AD pathogenesis.

Fibrinogen extravasates from the blood vessels into the brain parenchyma and is enzymatically converted to fibrin, which is deposited as diffuse perivascular leakage, along the blood vessel walls, and as dense deposits within amyloid plaques.^15, 18–20^ The conversion of fibrinogen to fibrin by thrombin exposes a cryptic epitope, γ377-395, which can bind to the CD11b i-domain of the complement receptor on microglia, promoting a pro-inflammatory response.^21^ Notably, fibrinogen knockouts or pharmacological depletion of fibrinogen using ancrod treatment in amyloid models reduces BBBd-induced neuroinflammation and amyloid pathology.^15, 18^ However, systemic depletion of fibrinogen carries significant risks, primarily the impairment of blood coagulation and increased risk of hemorrhage.^22^ To circumvent these risks, a mouse model lacking the pro-inflammatory CD11b receptor binding site on fibrin(ogen) (*Fgg*^γ390–396A^) that prevents CD11b receptor–mediated interactions with microglia without altering blood coagulation was engineered to determine the causal role of fibrin-CD11b interactions.^21, 23^ Notably, *Fgg*^γ390–396A^ reduces microglia-dependent neuroinflammation, synaptic dysfunction, and amyloid burden while improving cognitive and behavioral functions in 5xFAD mice^19, 24, 25^ Fibrin-targeting immunotherapy against γ377–395 has recently been developed with antibodies, including mouse 5B8 and humanized affinity-matured THN391 anti-fibrin antibodies, that specifically target fibrin without affecting coagulation.^26–28^ Treatment with the 5B8 anti-fibrin antibody (Kd = 15 nM) in 5xFAD mice reduced microglial activation, loss of cholinergic neurons, and inflammatory and oxidative stress responses.^27, 29^ Improved-affinity humanized THN391 antibodies (Kd = 0.1 nM) have recently been developed.^27^ THN391 antibody has recently entered Phase 1b clinical testing for safety and cognitive endpoints in patients with early AD. However, neither the clinical nor preclinical studies have defined the role of fibrin(ogen) on AD-related network dysfunction.

AD is increasingly recognized as a disorder of network dysfunction resulting in hyperactivity, disrupted sleep-circadian rhythms, and oscillatory slowing.^30^ Notably, BBBd and cerebral amyloid angiopathy (CAA) promote network hyperexcitability and epileptic activity in multiple neurological disorders.^31–35^ APP duplications or familial- and CAA-AD linked Aβ mutations, such as the Dutch and Iowa variants, causes early-onset AD with prominent CAA and seizures.^36–40^ Consistently, the BBBd subtype had enriched APP variants associated with CAA.^16^ In this context, it is important to note that fibrin(ogen) exhibits high-affinity binding to Aβ,^41^ with this affinity increasing 50-fold in CAA-linked Aβ familial AD mutations.^42^ Altogether, this suggests a strong mechanistic link between AD-related vascular pathology, fibrinogen, and network hyperexcitability. In addition to network hyperexcitability, AD patients also exhibited altered brain oscillatory rhythms, characterized by a prominent EEG slowing,^30, 43^ and sleep^44^ and circadian rhythm alterations^44–47^ often preceding cognitive decline.^44^ These aberrant patterns of brain network activity seem to contribute to AD-related pathogenesis, including Aβ and tau accumulation.^43, 46, 48–52^ However, the role of vascular dysfunction and neuroinflammation in AD-related brain network dysfunction, including hyperexcitability, altered brain oscillations, and sleep and circadian disturbances, as well as the efficacy of therapeutic targeting of these abnormalities with immunotherapy, have not been defined. Here, we used genetic (*Fgg*^γ390–396A^) or pharmacological antibody-based (5B8 and THN392, the murine version of the clinical-stage humanized fibrin-targeting antibody THN391) approaches to block fibrin-microglia interactions and assess the impact of vascular neuroinflammation on AD-related brain network dysfunction. We found that genetic (*Fgg*^γ390–396A^) manipulation or 5B8 and THN392 antibody treatment reduced 5xFAD-associated alterations in brain network activity, including hyperexcitability, altered brain oscillations, and sleep and circadian disturbances. We conclude that fibrin-induced microglia-mediated neuroinflammation promotes AD-related network dysfunction and that pharmacological targeting of the fibrin-microglia axis represents a potential therapy for network dysfunction in AD.

## RESULTS

### Fibrin(ogen) is deposited in the brain parenchyma in AD and related mouse models

To determine if fibrin(ogen) is deposited in the brain parenchyma of a variety of amyloid mouse models, including 5xFAD, *App*^SAA/SAA^, and *App*^NLG/NLG^:*APOE4* mice, and humans with AD, we stained brain sections for fibrin(ogen), Aβ, and microglia. As shown before in human and related mouse models,^15, 18–20, 53^, we identified three patterns of fibrin(ogen) in the parenchyma: strong fibrin(ogen) deposition within amyloid plaques, diffuse fibrin(ogen) around blood vessels indicative of perivascular leakage, and fibrin(ogen) localized around certain blood vessels (**Figure 1A**).

**Figure 1.**
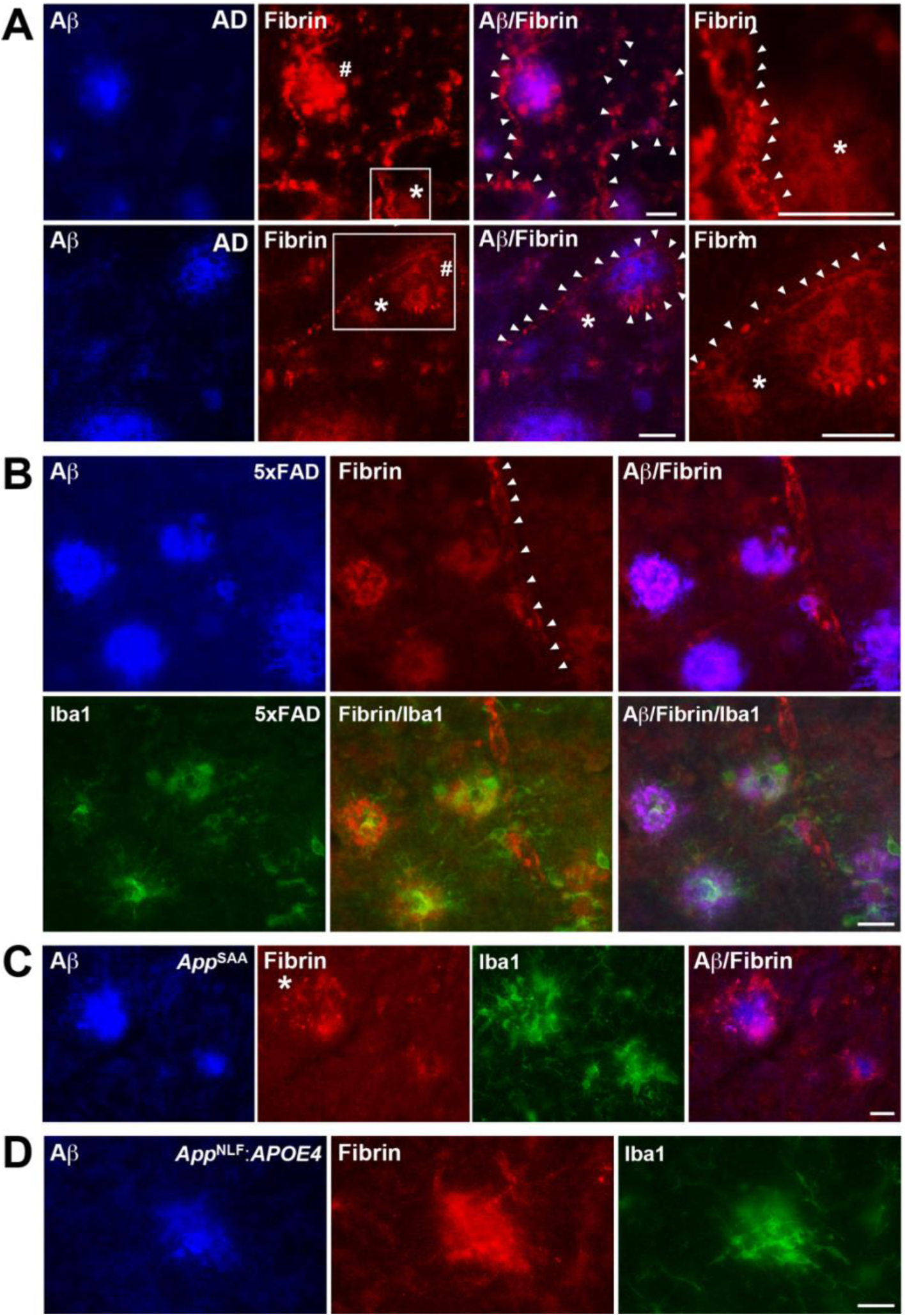
Fibrin(ogen) is deposited in the brain perivascular space in AD patients and 5xFAD, *App*^SAA/SAA^, *App*^NLG/NLG^:*APOE4* mouse models. Brain tissue from human AD (A) and from 5xFAD, *App*^SAA/SAA^, *App*^NLG/NLG^, and *App*^NLG/NLG^:APOE4 mice (B-D) were stained for fibrin(ogen) (red), Aβ (blue), and the microglia marker Iba1 (green) primary antibodies (scale bar, 25 mm). (**A**) In AD brain tissue, fibrin(ogen) deposition was found within amyloid plaques (#), diffusely around some of the blood vessels (*), and decorating blood vessels (arrowheads). Representative images are shown from n = 3 samples. (**B**) 10-month-old 5xFAD mice showed robust fibrin(ogen) deposition within and around amyloid plaques and decorating blood vessels (arrowheads). Reactive microglia (green) accumulated around amyloid and fibrin(ogen) deposits. Representative images are shown from n = 10 samples. (**C, D**) Relative to 5xFAD mice (B), 21-month-old *App*^NLG/NLG^ (C) and 15-month-old *App*^NLG/NLG^:APOE4 (D) mice showed more diffuse fibrin(ogen) deposition around amyloid plaques, with fibrin(ogen) forming an external halo around amyloid plaques. Reactive microglia cells (green) accumulated around amyloid and fibrin(ogen) deposits. Representative images are shown from n = 6–10 samples per genotype.

Next, we tested fibrin(ogen) deposition in the transgenic model 5xFAD and two novel humanized knock-in (KI) models of amyloidosis, *App*^SAA/SAA^ and *App*^NLG/NLG^:*APOE4* mice. Fibrin(ogen) has been shown to be deposited in transgenic APP mice,^15, 18, 19, 53^ but not in novel humanized knock-in mice without APP overexpression. We used aged mice with prominent amyloid deposition. As previously demonstrated,^19^ 10-month-old 5xFAD mice had robust fibrin(ogen) deposition around and within amyloid plaques (**Figure 1B**). Interestingly, we also found similar fibrin(ogen) deposition in 21-month-old *App*^SAA/SAA^ (**Figure 1C**) and 16-month-old *App*^NLG/NLG^:APOE4 mice (**Figure 1D**), indicating that humanized models also accumulate fibrin in their brains. Microglia marker Iba1 staining revealed that reactive microglia were highly concentrated around amyloid/fibrin(ogen) deposits in all mouse models (**Figures 1B**–**1D**). These results indicate that fibrinogen is extravasated from the blood vessels into the brain parenchyma and deposited as fibrin(ogen) in AD and related transgenic and KI AD models.

### Alterations in brain network hyperexcitability and brain oscillations are rescued by blocking fibrin-microglia interactions

AD results in brain network hyperexcitability and epileptic activity,^30, 54–58^ a feature consistently recapitulated across APP-Tg models^59–68^ and humanized *App*-KI mice.^60^ To determine whether fibrin-microglia interactions modulate AD-related epileptic activity, we conducted continuous wireless EEG/EMG recordings for seven days in 10–13-month-old control, 5xFAD, and 5xFAD:Fgg^γ390–396A^ mice (**Figure 2A**). Fggγ^390-396A^ mice express a fully coagulable fibrinogen that specifically lacks the CD11b receptor binding site. As previously described,^69^ we found prominent epileptiform activity in 5xFAD mice (**Figure 2B**). Epileptic activity was most prominent during the resting phase (day) of the circadian cycle (**Figure 2C, left**), when mice are typically inactive, indicating that AD-related epileptiform activity is strongly modulated by circadian rhythms. Consistently, epileptiform spikes were strongly modulated by locomotor activity levels across the circadian cycle in both 5xFAD and 5xFAD:*Fgg*^γ390–396A^ mice, with periods of low locomotor activity exhibiting higher epileptic spike rates and vice versa (**Figure 2C, right**). Notably, relative to 5xFAD mice, 5xFAD:Fgg^γ390–396A^ mice showed reduced epileptiform activity during both day and night, and across locomotor activity level (**Figure 2C**). To determine sex effects, we plotted epileptic spikes by locomotor activity level and sex. We found that male and female 5xFAD mice had similar activity-dependent epileptic spike rates, and that Fgg^γ390–396A^ reduced epileptic spikes in both sexes (**Figure S1A**). These results indicate that fibrin-microglia interactions promote AD-related network hyperexcitability.

**Figure 2.**
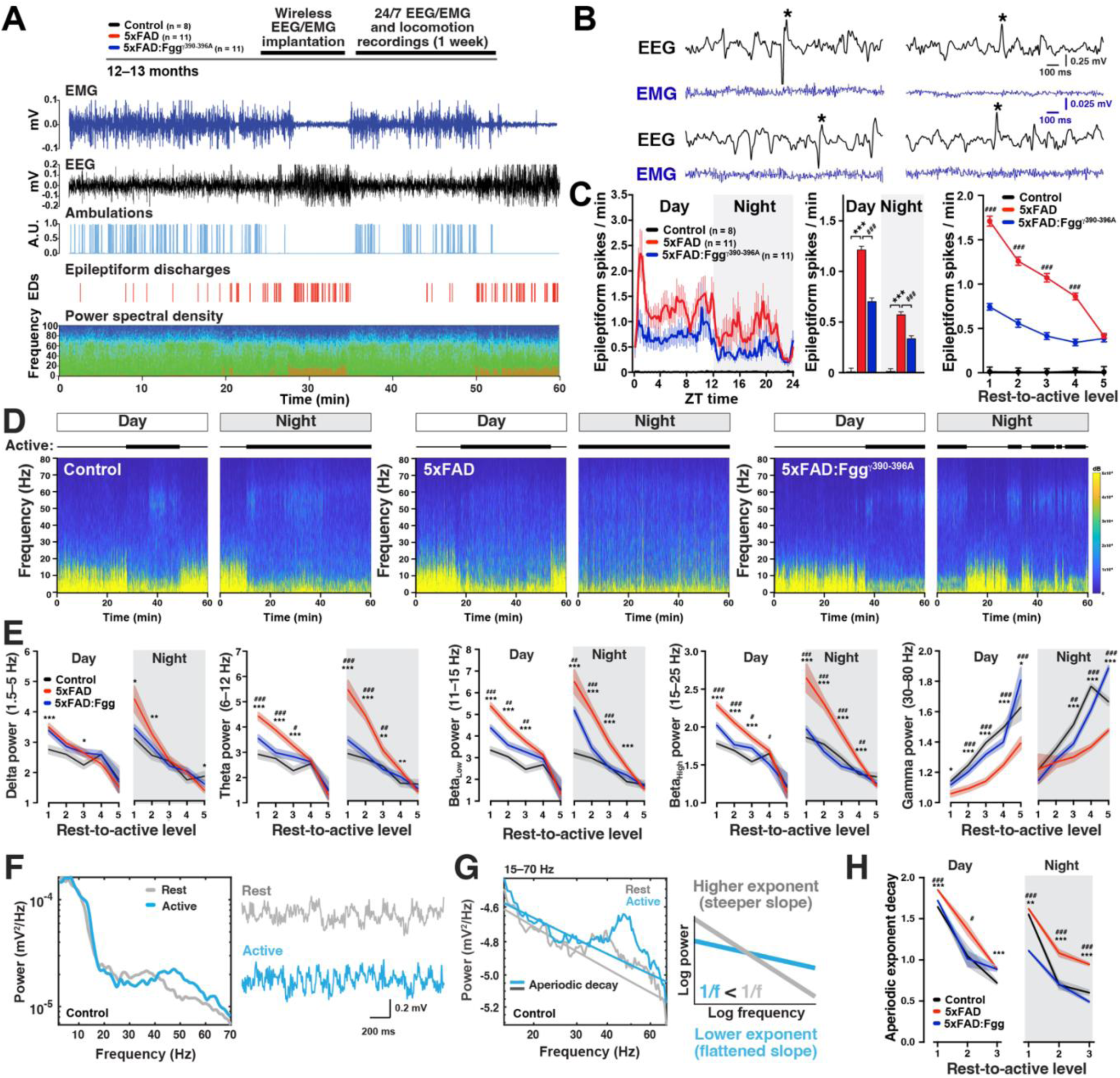
5xFAD-dependent alterations in network hyperexcitability and brain oscillations are ameliorated by genetically blocking fibrin–microglia interactions in 5xFAD:*Fgg*^γ390–396A^ mice. 10–13-month-old mixed-sex control (n = 8), 5xFAD (n = 11), and 5xFAD:*Fgg*^γ390–396A^ (n = 11) mice were implanted with a wireless EEG/EMG transmitter to continuously monitor cortical brain activity for 7 days. (**A**) Experimental timeline. Representative traces showing EMG (blue), EEG (black), ambulation (light blue), epileptiform spikes (red), and power spectral density over one hour of recording. Note epileptiform spikes emerge during periods of high delta and theta oscillatory power and low locomotion (5xFAD mouse). (**B**) Four representative EEG and EMG traces showing examples of subclinical epileptiform activity, characterized by abnormal epileptic discharges in the absence of EMG activity. (**C**) Averaged epileptiform spikes were plotted in 15-minute intervals along the 24-hour circadian cycle (left), for day versus night (middle), and by locomotor activity level (rest-to-active) (right). Relative to 5xFAD mice, 5xFAD:*Fgg*^γ390–396A^ mice had decreased epileptic activity across circadian and behavioral states. (**D**) Representative 1-hour spectrograms from day and night periods for control, 5xFAD, and 5xFAD:*Fgg*^γ390–396A^ mice. Note reduced gamma power (30-80 Hz) during active periods in 5xFAD mice. (**E**) Cortical brain oscillatory power of canonical oscillatory frequency bands, including delta (1.5–5 Hz), theta (6–12 Hz), low and high beta (11–15 Hz, 15–25 Hz), and gamma (30–80 Hz) per locomotor activity level (rest-to-active) for day and night (7 days of recordings). 5xFAD mice had increased power in low-frequency oscillations (delta, theta, and beta) and decreased power in gamma. Altered 5xFAD-induced oscillatory alterations were mostly restored in 5xFAD:*Fgg*^γ390–396A^ mice. (**F**) Power spectral density (PSD) and corresponding voltage EEG traces during resting and active periods in a control mouse. Note the increased power of gamma oscillations during active periods. (**G**) The aperiodic component of the PSD, computed over the 15–70 Hz range, was quantified as the exponent of the 1/f distribution. Resting periods had a higher exponent, corresponding with a steeper slope on the log–log frequency–power plot (higher decay), whereas active periods had a lower with a flatter log–log slope, reflecting reduced aperiodic decay over the 1/f distribution. (**H**) Aperiodic decay during day and night in control, 5xFAD, and 5xFAD:*Fgg*^γ390–396A^ mice over locomotor activity levels. Values are mean ± SEM; P values by Generalized Linear Mixed Model (GLMM) accounting for repeated measures, individual differences (random factor), and fixed factors (day/night for C; day/night and activity levels for E and H), and Bonferroni post hoc test for multiple comparisons. \**p* < 0.05, \*\**p* < 0.01, and \*\*\**p* < 0.001 for controls vs. 5xFAD; ^#^*p* < 0.05, ^##^*p* < 0.01, and ^###^*p* < 0.001 for 5xFAD vs. 5xFAD:*Fgg*^γ390–396A^

Brain oscillatory activity is closely related to cognitive functions and these brain rhythms are affected in AD.^30, 48, 70–76^ To determine whether fibrin-microglia interactions modulate AD-related oscillatory activity, we calculated the cortical brain oscillatory power of canonical frequency bands, including delta (1.5–5 Hz), theta (6–12 Hz), low and high beta (11–15 and 15–25 Hz), and gamma (30–80 Hz) per locomotor activity level (rest-to-active) for day and night for the 7-day recordings (**Figures 2D and 2E**). Similar to the EEG slowing described in AD patients,^77^ we found that 5xFAD mice had increased power in low-frequency oscillations, including delta, theta, and beta, at low locomotor activity levels (**Figure 2E**). This increased power of low frequency oscillations was particularly prominent during the resting periods of the night, indicating EEG slowing.^30, 78^ Conversely, high frequency oscillations, such as gamma, were reduced in 5xFAD mice, at both low 30-60 Hz and higher 60-80 Hz gamma frequencies (**Figure S1B**). Remarkably, most of the 5xFAD-dependent oscillatory abnormalities were reduced in 5xFAD:Fgg^γ390–396A^ mice, indicating that fibrin(ogen) via the γ377-395 epitope is a key driver of AD-related EEG slowing.

The aperiodic decay, or slope, of the power spectral density (PSD), which is the non-oscillatory component of the power spectrum, has been recently identified as a potential biomarker of neurophysiological processes in humans (**Figure 2F**).^77, 79–85^ Higher exponents or steeper slopes have been linked to resting brain states, like sleep^82^ or propofol-induced anesthesia^81^, whereas lower exponents or flatter slopes are associated with active brain states (**Figure 2G**). Consistently, we found an inverse relationship between aperiodic decay and locomotor activity levels (**Figure 2H**), indicating a flatter aperiodic slope during active locomotion. Notably, as low-frequency oscillations are elevated a significant decrease in gamma power is observed in 5xFAD mice (**Figure 2E and 2H**). Relative to wild-type controls, we found that 5xFAD mice exhibited a steeper aperiodic slope across locomotor activity levels during both day and night, which was decreased by Fgg^γ390–396A^ expression in 5xFAD mice, indicating that fibrin(ogen) drives a shift toward abnormally elevated resting-state activity in 5xFAD mice. Altogether, these results indicate that fibrin-microglia interactions in 5xFAD mice drives epileptiform activity, brain slowing, and increased resting states.

### Alterations in sleep-circadian brain and behavioral rhythms in 5xFAD mice are rescued by blocking fibrin-microglia interactions

Sleep patterns and circadian rhythms are disrupted in AD and represent an early indication of dementia.^44, 45^ To determine if sleep architecture is altered in 5xFAD mice, we measured the time spent in each sleep stage and the number of sleep-stage transitions throughout the seven days of recording. During the day (resting phase), 5xFAD mice exhibited robust increases in transitions from Wake to Sleep (**Figure 3A**) and in total Sleep time (**Figure S2A**). Conversely, during the night (active phase), 5xFAD mice exhibited decreased transitions from Wake to Sleep (**Figure S2B**) and reduced Sleep time (**Figure S2A**). These results indicate that 5xFAD mice exhibit excessive daytime sleepiness (hypersomnolence) and nocturnal restlessness (insomnia),^86–88^ Beyond total sleep stage durations, 5xFAD mice displayed marked alterations in sleep architecture and increased fragmentation. This was characterized by a higher bidirectional frequency of transitions between REM and NREM (**Figure 3A**), an increase in REM bout duration alongside a reduction in NREM bout duration (**Figure 3B**), and a lower threshold for REM entry and exit—evidenced by increased Wake-to-REM (entry) and REM-to-Wake (exit) transitions (**Figure S2C**). These findings are consistent with REM intrusion/arousal instability phenotypes observed in AD patients with excessive daytime sleepiness and uncontrollable bouts of REM sleep (arousal instability) at any time, which has been linked deficits in the orexin (hypocretin) system.^89, 90^ Our results indicate significantly disrupted sleep patterns in the 5xFAD mice that were largely ameliorated in the 5xFAD:*Fgg*^γ390–396A^ mice, suggesting that fibrin(ogen) contributes to AD-related sleep alterations.

**Figure 3.**
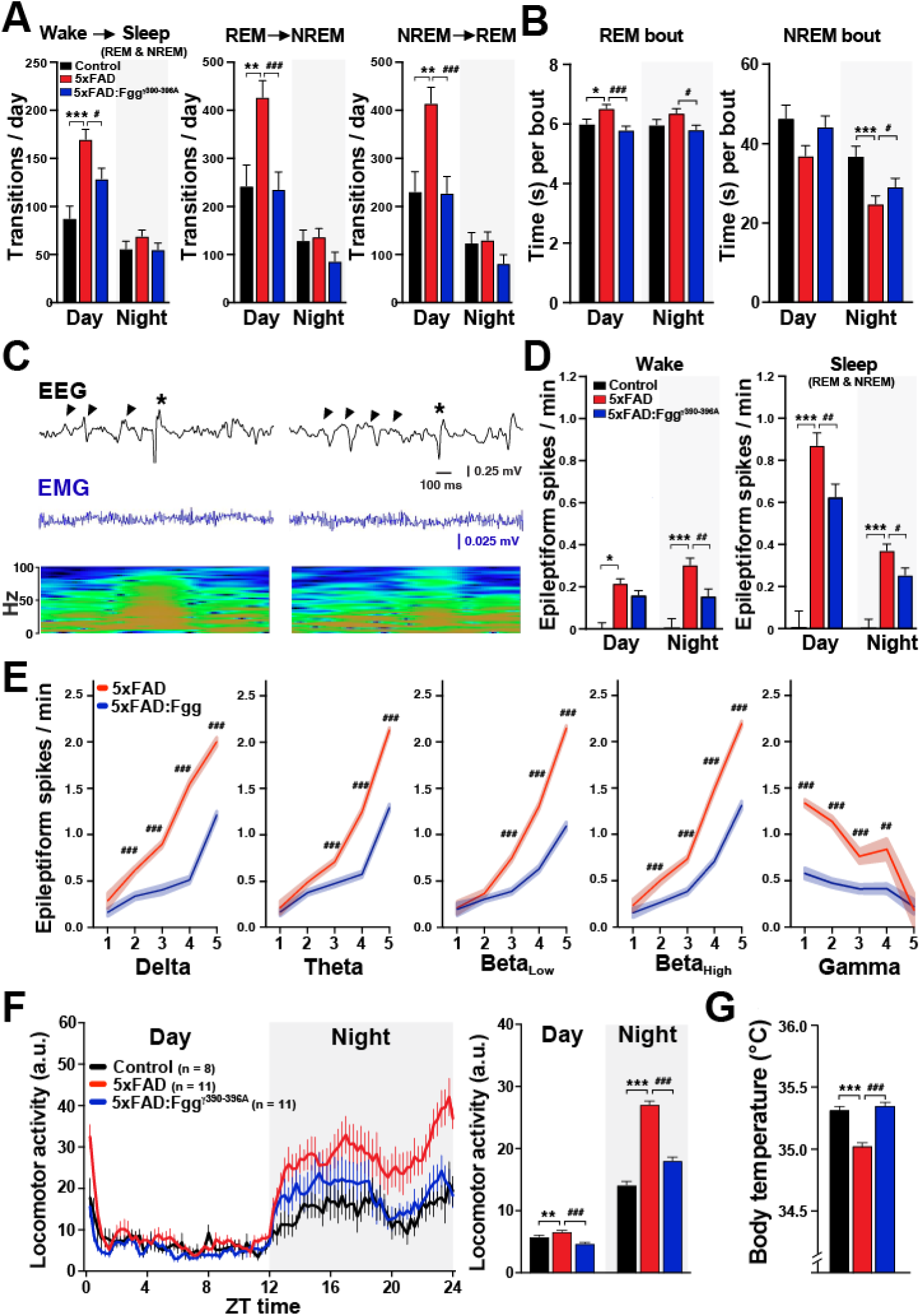
5xFAD-dependent alterations in sleep patterns, circadian activity, and hyperactivity are reduced by blocking fibrin–microglia interactions in 5xFAD:*Fgg*^γ390–396A^ mice. 10–13-month-old mixed-sex control (n = 8), 5xFAD (n = 10), and 5xFAD:*Fgg*^γ390–396A^ (n = 10) mice were implanted with a wireless EEG/EMG transmitter to continuously monitor sleep activity, locomotion, and temperature for 7 days under home cage conditions. (**A**) Average number of sleep stage transitions during day and night from Wake to Sleep (REM and NREM) (left), from REM to NREM (center), and from NREM to REM (right). 5xFAD mice had increased transitions to Sleep and between sleep stages, which was ameliorated in 5xFAD:*Fgg*^γ390–396A^ mice. (**B**) REM and NREM bout duration. 5xFAD mice had increased REM bout duration, but decreased NREM bout duration, which were ameliorated in 5xFAD:*Fgg*^γ390–396A^ mice. (**C**) Epileptic spike EEG traces (black), EMG signal (blue), and spectrograms showing subclinical epileptiform activity (asterisks) during sleep periods with slow-wave oscillations (arrowheads) and increased delta power. (**D**). Sleep stages and epileptic activity. Average number of epileptiform spikes in Wake (left) and Sleep (REM and NREM, right) for the 7 days of recordings. Epileptic activity was prominent during Sleep and the resting period of the circadian cycle (day). 5xFAD:*Fgg*^γ390–396A^ mice had reduced 5xFAD-dependent epileptic activity. (**E**) Epileptiform activity was associated with slow-frequency oscillations (delta, theta, beta) and negatively associated with fast-frequency oscillations (gamma) in 5xFAD mice, which was ameliorated in 5xFAD:*Fgg*^γ390–396A^ mice. (**F**) Locomotion recorded across 7 days in 15-minute bins along the 24 h circadian cycle (left) or averaged for the day and night (right). Relative to controls, 5xFAD mice had increased locomotion particularly during the night, which was reduced in the 5xFAD:*Fgg*^γ390–396A^ mice. (**G**) Temperature (24 hr, day and night) recorded across 7 days in relation to locomotor activity level. Relative to controls, 5xFAD mice had decreased body temperature, which was ameliorated in 5xFAD:*Fgg*^γ390–396A^ mice. Values are mean ± SEM; P values by Generalized Linear Mixed Model (GLMM) accounting for repeated measures and mouse (random effect), fixed factors (day/night for A, B, D, F; power levels for E; activity levels for G) and Bonferroni post hoc test for multiple comparisons; \**p* < 0.05, \*\**p* < 0.01, and \*\*\**p* < 0.001 for Controls vs. 5xFAD; ^#^*p* < 0.05, ^##^*p* < 0.01, and ^###^*p* < 0.001 for 5xFAD vs. 5xFAD:*Fgg*^γ390–396A^

Sleep stages modulate AD-related epileptic activity in humans and related mouse models.^91, 92^ To assess this relationship, we determined the number of epileptiform spikes per sleep stage. Consistent with human^91,92^ and other APP mouse models (APP/PS1),^93^ epileptic activity was more prominent during the resting phase of the circadian cycle and sleep stages (**Figures 3C and 3D**), than during wakefulness, in 5xFAD mice, indicating that a resting brain stage promotes AD-related network hypersynchrony. Blocking fibrin-microglia interactions in 5xFAD:*Fgg*^γ390–396A^ mice reduced epileptic activity during both Wake and Sleep stages. We noticed that epileptiform spikes emerged during periods of high delta power and low locomotion (**Figure 2A**). To demonstrate this relationship, we correlated the levels of epileptic activity with slow- and fast-frequency oscillatory bands. We found that epileptiform activity was positively associated with slow-frequency oscillations (delta, theta, beta) and negatively associated with fast-frequency oscillations (gamma) (**Figure 3E**). Notably, the epileptiform spikes were preceded by slow-wave, large-amplitude, and low-frequency EEG activity (**Figure 3C**). Consistent with oscillatory changes in **Figure 2**, 5xFAD:*Fgg*^γ390–396A^ mice showed a reduction of 5xFAD-induced epileptic activity in an oscillatory-dependent manner.

Sleep abnormalities in AD are linked to the disruption of behavioral circadian activity patterns.^46, 94^ To determine whether AD-related circadian activity patterns are present in 5xFAD mice and modulated by fibrin-microglia interactions, we continuously monitored locomotion and body temperature under home cage conditions for seven days in 10–13-month-old control, 5xFAD, and 5xFAD:Fgg^γ390–396A^ mice using wireless subcutaneous transmitters. Consistent with the nocturnal restlessness phenotype described above, 5xFAD mice showed increased locomotion relative to controls, particularly during the active night cycle (**Figure 3F**). This hyperlocomotion was nearly normalized in 5xFAD:*Fgg*^γ390–396A^ mice, indicating that fibrin-microglia interactions promote circadian-dependent overnight hyperlocomotion in 5xFAD mice. Notably, we also found that the reduced body temperature observed in 5xFAD mice was rescued in 5xFAD:*Fgg*^γ390–396A^ mice (**Figure 3G**). Altogether, these findings demonstrate that 5xFAD mice exhibit multifaceted deficits in sleep-circadian homeostasis that are mostly driven by fibrin-microglia interactions and are largely mitigated by genetically inhibiting fibrin-CD11b binding.

### Anxiety-like behavior in 5xFAD mice is rescued by blocking fibrin-microglia interactions

Anxiety is one of the most common neuropsychiatric symptoms in AD, affecting nearly 40% of patients,^95^ and is often observed preceding the diagnosis of dementia in amyloid-positive individuals,^96^ predicting the conversion from MCI to AD.^97^ We have previously shown that *Fgg*^γ390–396A^ protects against cognitive impairment and spontaneous behavioral alterations in 5xFAD mice,^19, 25^ but its effects on anxiety have not yet been defined. To test this, we performed the open field test in 7–10-month-old control, 5xFAD, and 5xFAD:Fgg^γ390–396A^ mice during the daytime. Consistent with the overnight hyperlocomotion (**Figure 3F**), we found robust hyperlocomotion in 5xFAD mice when exploring a novel environment (**Figure 4A**). Notably, this hyperactivity was mainly driven by impairments in habituation–a basic form of contextual learning^98, 99^– with 5xFAD mice exhibiting aberrant and continuous exploratory behavior that did not decay over the 60-min open field trial (**Figure 4B**). These alterations were greatly normalized in 5xFAD:*Fgg*^γ390–396A^ mice (**Figures 4A and 4B**). We found similar effects across both sexes regarding the severity of 5xFAD-induced phenotype and the protection conferred by *Fgg*^γ390–396A^ (**Figure 4A, right**). Hyperlocomotion and habituation deficits in the open field have been observed in multiple APP-Tg and App-KI mice and are attributed to processing deficits of contextual information.^98–102^ To assess anxiety-like behavior, we quantified the time spent in the exposed center zone versus the secure periphery (edges) of the open field arena. We found that 5xFAD mice spent more time in the periphery and less time in the center (**Figure 4C**), suggesting an increased anxiety-like phenotype. This deficit was largely reversed in 5xFAD:*Fgg*^γ390–396A^ mice, suggesting a fibrin-microglial-dependent anxiety mechanism in 5xFAD mice.

**Figure 4.**
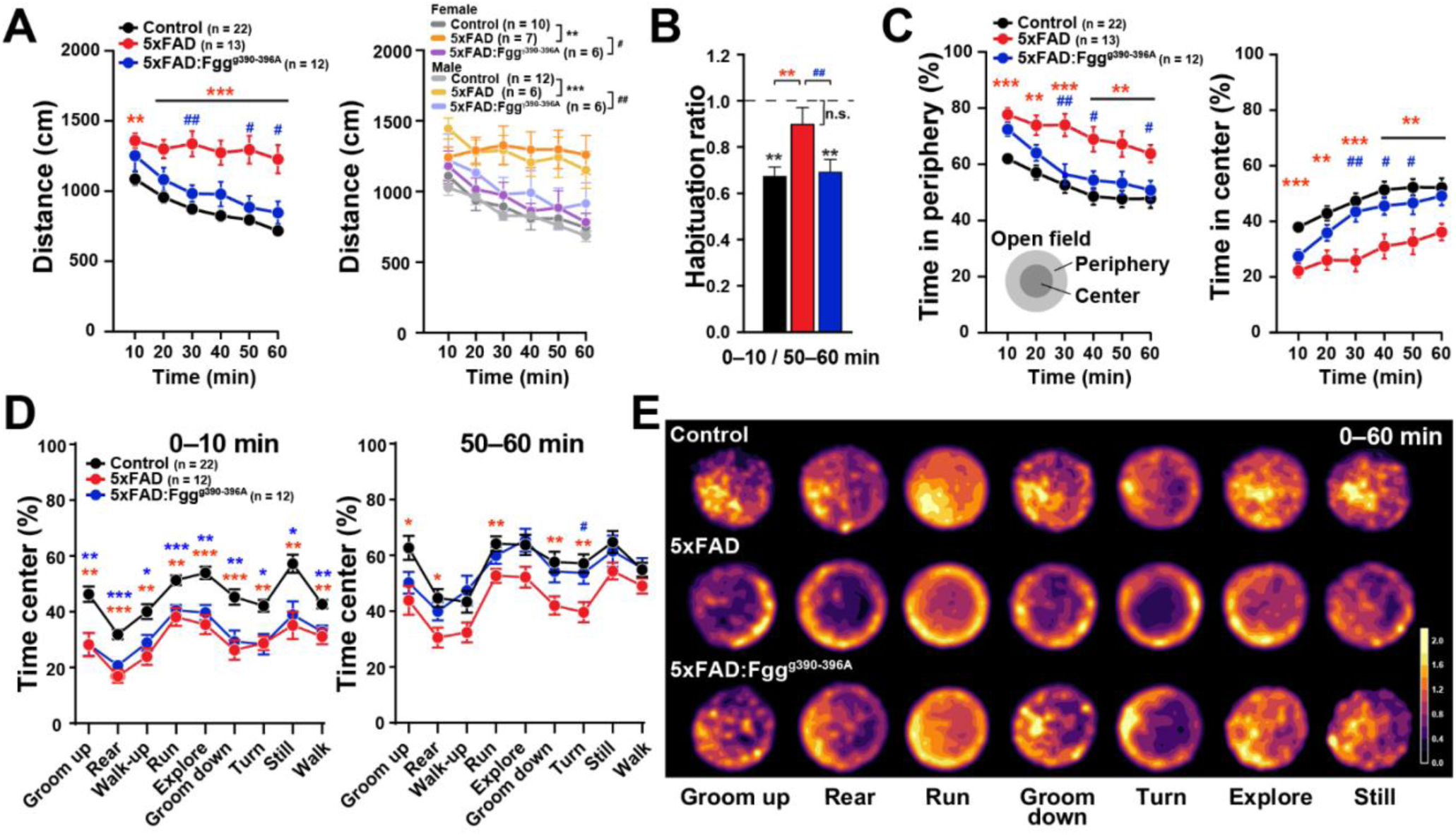
Machine learning open field spatial analysis reveals 5xFAD behavioral alterations in anxiety and hyperactivity are ameliorated by blocking fibrin-microglia interactions. 7–10-month-old mixed-sex control (n = 22), 5xFAD (n = 13), and 5xFAD:*Fgg*^γ390–396A^ (n = 12) mice were tested in the open field for 60 minutes and assessed with the machine learning VAME platform.^25^ (**A**) Distance traveled in 10-min intervals during the 60-min open field test, presented for both sexes combined (left) and disaggregated by sex (right). Both female and male 5xFAD mice (n = 7 females and 6 males) showed hyperactivity, which was ameliorated in 5xFAD:*Fgg*^γ390–396A^ mice (n = 6 females and 6 males). (**B**) Habituation index for the first and last 10-min intervals of the open field test (0–10 min / 50–60 min). 5xFAD mice failed to habituate, denoting a contextual learning deficit, which was rescued in 5xFAD:*Fgg*^γ390–396A^ mice. \*\**p* < 0.01 and n.s. (black) by t test relative to 1 (no habituation). (**C**) Time spent in the periphery (left) and in the center (right). 5xFAD mice exhibited a decrease in time spent in the center, denoting an anxiety-like phenotype, which was greatly attenuated in 5xFAD:*Fgg*^γ390–396A^ mice. (**D**) Behavioral communities in the center (% time) from VAME spatial analysis across the first (0–10 min; left) and last (50–60 min; right) ten open field minutes. VAME communities revealed that 5xFAD:*Fgg*^γ390–396A^ behaviors did not differ from WT controls across communities during the last 10 min of open field. (**E**) Time spent heatmaps for VAME communities indicating usage across 0–60 min of open field Values are mean ± SEM; P values by repeated two-way ANOVA with Tukey’s test for multiple comparisons (A left, C, D), and by one-way ANOVA with Holm-Sidak’s test for multiple comparisons (A, right); \**p* < 0.05, \*\**p* < 0.01, and \*\*\**p* < 0.001 (red asterisk) for Controls vs. 5xFAD, (blue asterisk) for Control vs. 5xFAD:*Fgg*^γ390–396A^; ^#^*p* < 0.05, ^##^*p* < 0.01, and ^###^*p* < 0.001 for 5xFAD vs. 5xFAD:*Fgg*^γ390–396A^

To better understand the behaviors associated with center-related anxiety, we performed a spatial analysis of 30 behavioral motifs–short, unique behavioral sequences identified using the machine learning platform VAME.^25, 103^ We found that 28 out of 30 behavioral motifs (**Figure S3A**) and all nine behavioral communities (**Figure S3B**) showed a significant decrease in the percentage of time spent in the center in 5xFAD mice, compared to controls. Notably, when we assessed the first (0–10 min) and the last (50–60 min) ten-min intervals of the open field trial, relative to controls, all nine behavioral communities in both 5xFAD and 5xFAD:*Fgg*^γ390–396A^ mice showed decreased time within the center during the first 10 minutes relative to controls (**Figure 4D**, left). However, by the last 10 minutes of the OF trial, 5xFAD mice continued to spend significantly less time in the center of the arena compared to controls (**Figure 4D**, right), whereas the behavior of 5xFAD:*Fgg*^γ390–396A^ mice was largely normalized. Interestingly, the Rear and Explore behavioral communities showed the most prominent and meaningful rescue (**Figure S3B**), as visualized by the spatial heatmaps (**Figure 4E**). Thus, our results suggest that *Fgg*^γ390–396A^ protects against habituation deficits and anxiety alterations in 5xFAD mice.

### Treatment with the anti-fibrin antibody 5B8 mitigates alterations in circadian activity, network hyperexcitability, and oscillations in 5xFAD mice

5B8 is a monoclonal antibody that targets the cryptic fibrin epitope γ377-395, selectively inhibiting fibrin-induced inflammation and oxidative stress without interfering with clotting.^29^ 5B8 antibody treatment inhibits microglia activation and prevents cholinergic neurodegeneration in 5xFAD mice.^29^ However, the effects of 5B8 immunotherapy on brain network hyperexcitability and oscillations have not yet been determined. To address this question, we dosed (i.p.) 12–15-month-old mice every 48 hours for 1 week with saline (baseline recording), followed by 2 weeks of dosing every 48 hours (8 doses) with 30 mg/kg 5B8 (5xFAD-5B8) or IgG2b (5xFAD-IgG2b) antibody while EEG/EMG and locomotor activity were continuously recorded over the 3-week treatment period in their home cage (**Figure 5A**). 5B8 spatially correlates with fibrin-rich areas in 5xFAD brains after i.p. administration demonstrating target engagement.^29^ Locomotor activity and epileptiform spikes were normalized to the initial 7-days saline dosing period to assess longitudinal progression relative to baseline (Week 1). Notably, 5xFAD-IgG2b mice exhibited time-dependent increases in hyperlocomotion over the 3-week period, which were prevented in 5xFAD-5B8 mice (**Figure 5B**), indicating 5B8 efficacy after one or two weeks of treatment. To determine the effects on circadian activity, we assessed locomotion in 15-minute bins across the 24-h circadian cycle. 5B8 treatment significantly reduced overnight hyperlocomotion with no effects on diurnal activity (**Figure 5C**), counteracting the susceptibility of 5xFAD mice to overnight hyperlocomotion (**Figure 3F**).

**Figure 5.**
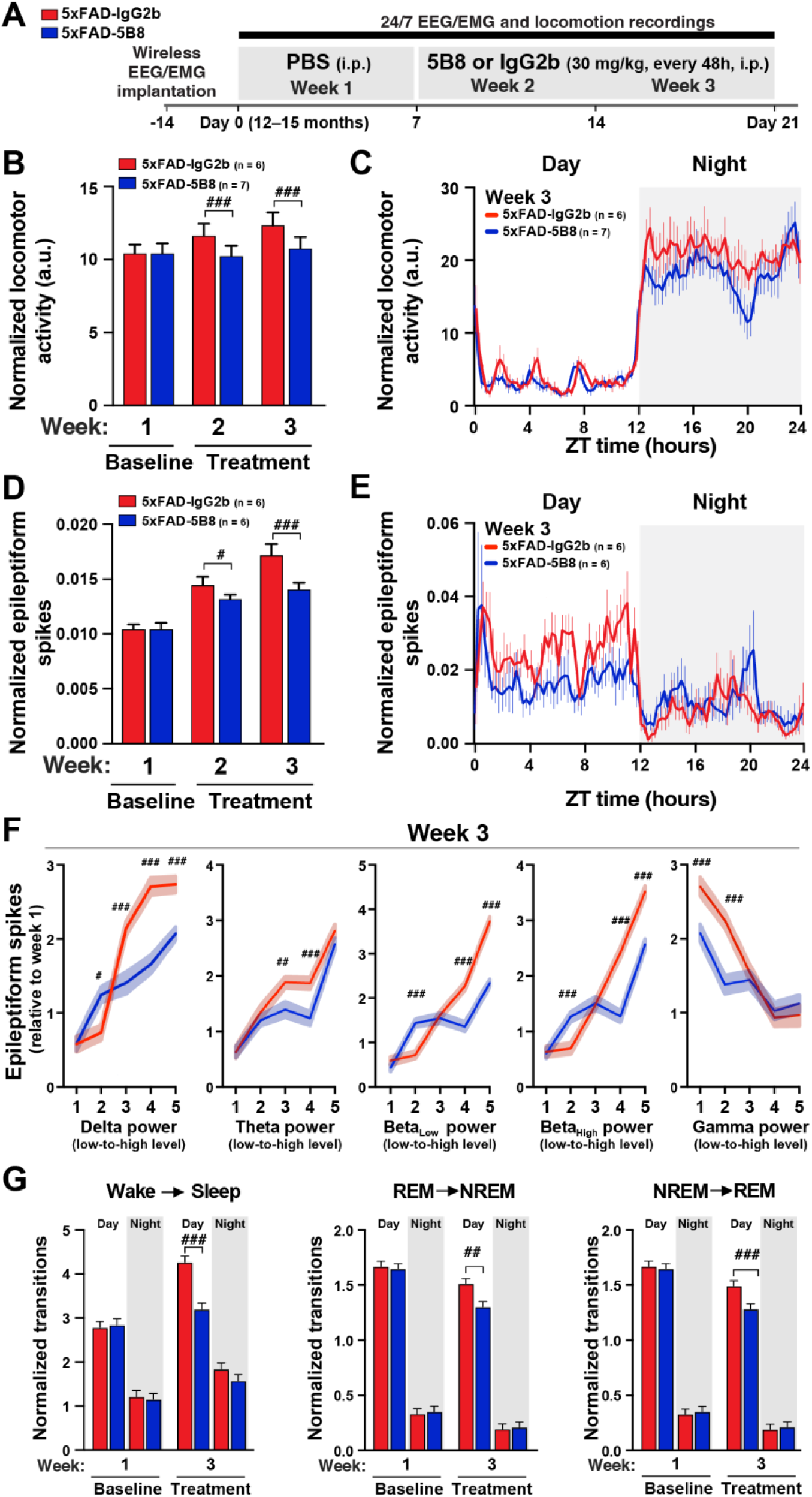
Acute (2 weeks) treatment with the anti-fibrin antibody 5B8 mitigates alterations in circadian activity, network hyperexcitability, and oscillations in 5xFAD mice. 12–15-month-old mixed-sex mice were dosed (i.p.) every 48 hours for 1 week with saline (baseline) followed by 2 weeks of dosing (8 total doses) every other day at 30 mg/kg with the 5B8 antibody (n = 6) or with IgG2b control antibody (n = 6) while EEG/EMG and locomotor activity were recorded. (**A**) Diagram of the experimental design. (**B-C**) Locomotion recorded across week 1 (baseline; saline) and weeks 2 and 3 (treatment; 5B8 or IgG2b) (B) or in 15-minute bins along the 24 hour circadian cycle (left) for week 3 (C). Values are normalized to week 1 (baseline; saline). 5B8 treatment decreased hyperactivity in 5xFAD mice. (**D-E**) Epileptiform activity recorded across week 1 (baseline; saline) and weeks 2 and 3 (5B8 or IgG2b) (**D**) or in 15-minute bins along the 24 hour circadian cycle (left) for week 3 (E). Values are normalized to week 1 (baseline, saline). 5B8 treatment decreased epileptic spikes in 5xFAD mice. (**F**) Epileptic activity per oscillatory power level (low-to-high) for the 7 days of recordings. Epileptiform activity was associated with slow-frequency oscillations (delta, theta, beta) and negatively associated with fast-frequency oscillations (gamma) in 5xFAD mice. 5B8 treatment reduced epileptic activity particularly during periods with high power of slow-frequency oscillations and low power of fast-frequency oscillations in 5xFAD mice. (**G**) Average number of sleep stage transitions from Wake to Sleep (REM and NREM) (left), from REM to NREM (center), and from NREM to REM (right). 5B8 treatment reduced transitions to Sleep and between sleep stages. Values are mean ± SEM; P values by Univariate General Linear Model (UGLM) accounting for repeated measures, fixed factors (treatment by weeks for B and D; power levels for F) and Bonferroni post hoc test for multiple comparisons; ^#^*p* < 0.05, ^##^*p* < 0.01, and ^###^*p* < 0.001 for 5xFAD-IgG2b vs. 5xFAD-5B8.

We next determined the effects of 5B8 treatment on 5xFAD-induced epileptic activity. 5xFAD-IgG2b mice exhibited marked time-dependent increases in epileptic activity over time, which was ameliorated by 5B8 treatment in 5xFAD-5B8 mice (**Figure 5D**). Assessment of epileptic activity across the 24-h circadian cycle revealed that 5B8 treatment predominantly reduced epileptic activity during the day when mice are less active (**Figure 5E**), counteracting the diurnal susceptibility of 5xFAD mice to network hyperexcitability during the resting phase of the circadian cycle (**Figure 2C**). When assessing the relationships between epileptic activity and power of frequency bands, we found that 5B8 treatment reduced 5xFAD-induced epileptiform activity when it was more prominent, specifically during periods of high power in low-frequency bands (delta, theta, and beta) and low power across high-frequency bands (gamma) (**Figures 5F and S4A**) and during Sleep (**Figures S4B**), counteracting 5xFAD-dependent effects (**Figure 3E**). We also found that 5B8 treatment reduced transitions from Wake to Sleep and between sleep stages (**Figure 5G**) similar to the rescue observed in the 5xFAD:*Fgg*^γ390–396A^ mice (**Figure 3A**). Altogether, these results indicate that one or two weeks of 5B8 treatment is sufficient to counteract 5xFAD-dependent effects on circadian locomotion, network hypersynchrony, and sleep transitions.

### Treatment with anti-fibrin antibody THN392 mitigates alterations in brain network and behavior in 5xFAD mice

We evaluated the chronic effects of THN392, the murine version of the clinical-stage human fibrin-targeting antibody THN391.^27^ Both THN391 and THN392 are affinity-matured antibodies that target the fibrin γ377-395 epitope. THN391 and THN392 exhibit a 150-fold greater affinity for γ377-395 (Kd < 0.1 nM) than the parent antibody 5B8 (Kd ∼15 nM). THN391 is currently in clinical trials for early AD.^27^ The murine version, THN392, incorporates murine constant regions to avoid the development of anti-drug antibodies. 7–8-month-old 5xFAD mice were dosed biweekly (i.p.) for 13 weeks (25 doses) with 30mg/kg THN392 (5xFAD-THN392) or IgG2b control (5xFAD-IgG2b) antibody (**Figure 6A**). THN392 spatially correlates with fibrin-rich areas in 5xFAD brains after i.p. administration demonstrating target engagement.^27^ After 10 weeks of treatment, mice were implanted with EEG/EMG transmitters and continuously recorded for seven days in their home cage. Twelve weeks of THN392 treatment prominently reduced nighttime hyperlocomotion in 5xFAD mice (**Figure 6B**), effectively reversing the susceptibility of 5xFAD mice to overnight hyperlocomotion (**Figure 3F**). THN392 also prominently reduced epileptic activity, particularly during daytime, when 5xFAD mice have the most epileptic activity (**Figure 6C**). Notably, *Fgg*^γ390–396A^, 5B8, and THN392 interventions consistently demonstrated greater efficacy against hyperlocomotion and epileptiform activity during periods when these deficits are most pronounced in 5xFAD mice–nocturnal and diurnal, respectively. Relative to 5xFAD-IgG2b controls, treatment with THN392 exhibited superior efficacy compared to 5B8, likely due to its higher affinity (Kd < 0.7 vs ∼100 nM) and the extended duration of the treatment (2 vs 12 weeks).

**Figure 6.**
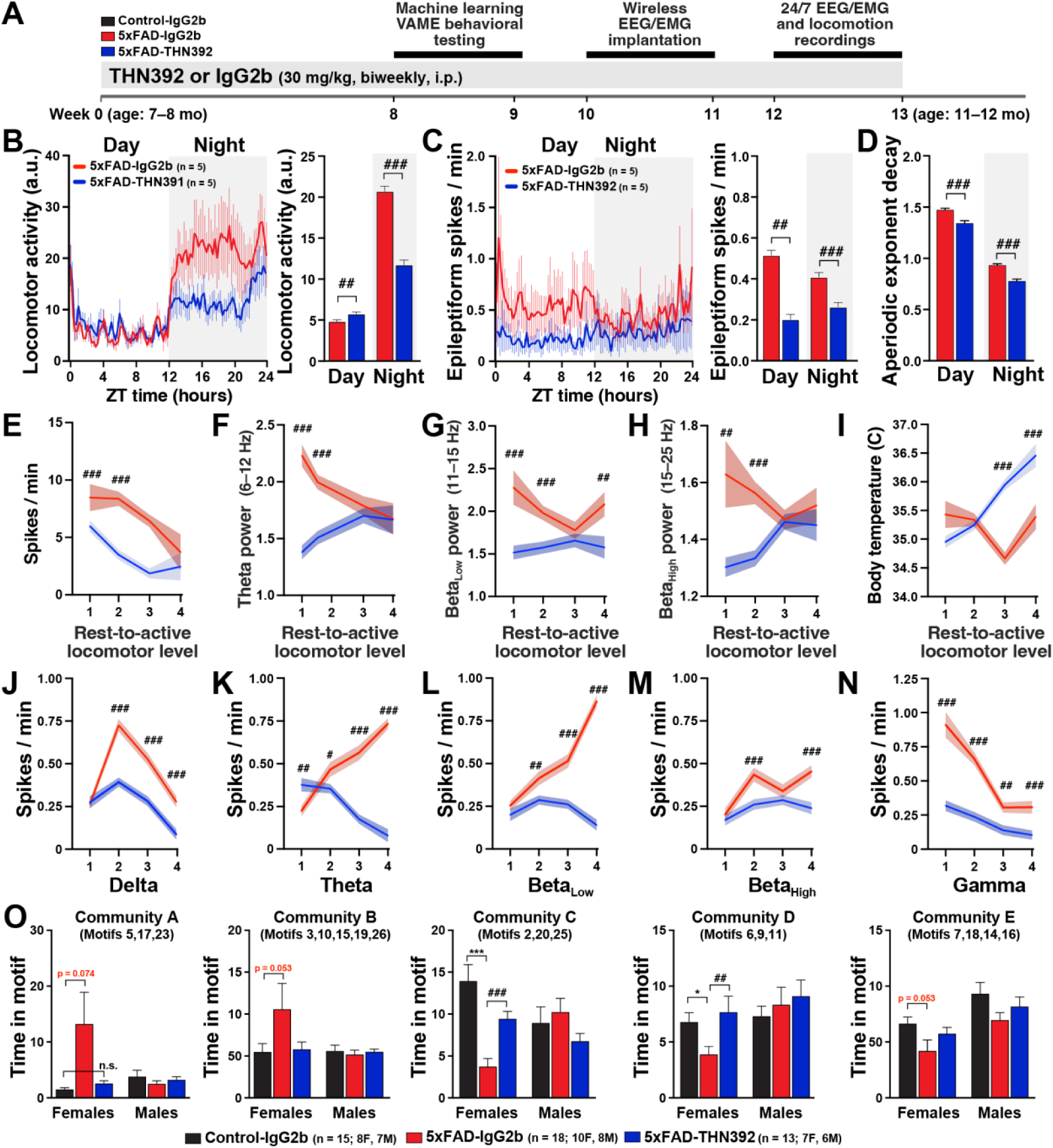
Treatment with anti-fibrin antibody THN392 mitigates alterations in circadian activity, network hyperexcitability, oscillations, and spontaneous behavior in 5xFAD mice. 7–8-month-old mixed-sex mice were biweekly treated (i.p.) for 13 weeks (25 total doses) at 30 mg/kg with the THN392 (n = 5) or IgG2b (n = 5) antibody. Behavior and 7-day cortical EEG/EMG activity were recorded at 8–9 and 10–11 weeks post-treatments, respectively. (**A**) Diagram of the experimental design. (**B-C**) Locomotion (B) and epileptiform activity (C) recorded and averaged across 7 days in 15-minute bins along the 24 hour circadian cycle (left) or averaged for the day and night (right). THN392 treatment reduced hyperlocomotion and epileptic activity in 5xFAD mice. (**D**) Aperiodic decay during day and night in IgG2b- and THN392-treated 5xFAD mice. THN392 treatment reduced the aperiodic decay in 5xFAD mice, counteracting 5xFAD-dependent effect (Figure 1H) and indicating a shift to more active brain states. (**E**) Epileptiform activity per locomotor activity level (rest-to-active). THN392 treatment reduced epileptic activity, particularly during resting periods with low levels of locomotor activity. (**F-H**) Cortical brain low-frequency oscillatory power of theta (F), beta (G, H) per locomotor activity level (rest-to-active). THN392 treatment reduced the power of low-frequency oscillations, counteracting the 5xFAD-dependent effects (Figure 1E). (**I**) Body temperature per locomotor activity level (rest-to-active). THN392 treatment increased body temperature, counteracting the 5xFAD-dependent effect (Figure 3G). (**J-N**) Epileptic activity per oscillatory power level (low-to-high). THN392 treatment decreased epileptic activity particularly during periods with high power of slow-frequency oscillations (delta, theta, and beta) and low power of fast-frequency oscillations in 5xFAD mice. (**O**) 7–8-month-old 5xFAD mice were biweekly treated (i.p.) for 8 weeks (16 doses) with the THN392 (n = 13; 7 females, 6 males) or IgG2b (n = 18; 10 females, 8 males) antibody and control mice with IgG2b (n = 15; 8 females, 7 males). Spontaneous behavior was assessed in the open field with the computer vision machine learning VAME platform. Relative to female control-IgG2b mice, female 5xFAD-IgG2b mice had robust behavioral alterations in the behavioral communities C and D, which were ameliorated in 5xFAD-THN392 mice. Values are mean ± SEM; P values by Generalized Linear Mixed Model (GLMM) accounting for repeated measures and mouse (random effect), fixed factors (day/night for B, C, D; activity levels for E-I; power levels for J-N) and Bonferroni post hoc test for multiple comparisons, and by one-way ANOVA and Bonferroni test for multiple comparisons (O); \**p* < 0.05, \*\**p* < 0.01, and \*\*\**p* < 0.001 for Controls vs. 5xFAD; ^#^*p* < 0.05, ^##^*p* < 0.01, and ^###^*p* < 0.001 for 5xFAD-IgG2b vs. 5xFAD-THN392

Since epileptic activity in 5xFAD mice robustly emerges during periods of aberrant increases in low-frequency power, including theta and beta oscillations, during resting states (**Figure 2E and 3E**), we assessed whether the antiepileptic effect of THN392 was associated with the amelioration of these oscillations. Indeed, we found that THN392 treatment attenuated 5xFAD-induced alterations in theta and beta power, particularly during resting states (**Figures 6F**–**6H**), while also reducing the aperiodic decay exponent (**Figure 6D**). Interestingly, THN392 also increased body temperature (**Figure 6I**), reversing the hypothermia observed in 5xFAD mice (**Figure 3G**). Consistent with the *Fgg*^γ390–396A^ and 5B8 manipulations, THN392 treatment reduced 5xFAD-induced epileptiform activity, predominantly during periods of low-frequency power (**Figures 6J**–**6N**), counteracting the 5xFAD-dependent effects (**Figure 3E**). These results suggest that the THN392 antibody may reverse the hyperexcitable resting state in 5xFAD mice by reducing aberrant low-frequency oscillations, thereby reducing overall hyperexcitability and shifting the brain into a more active brain state.

To determine the behavioral effects of THN392 on spontaneous behavior in the open field (60 min), we behaviorally tested 9–11-month-old Control-IgG2b, 5xFAD-THN392, and 5xFAD-IgG2b after 8 weeks of treatment using the computer vision machine learning VAME platform.^25^ Interestingly, we found that female, but not male, 5xFAD mice had behavioral abnormalities in VAME-defined behavioral communities C and D (**Figure 6O**). These deficits were associated with impaired gait and hunchback posture (kyphosis) observed in aged female, but not in male, 5xFAD mice, which were identified using VAME. Remarkably, nearly all of these motor deficits were restored in 5xFAD-THN392 mice, indicating restoration of gait and posture by THN392 treatment.

## DISCUSSION

We investigated the role of BBBd and fibrin(ogen)-induced neuroinflammation in AD-related brain network dysfunction, including network hyperexcitability, oscillatory alterations, sleep disruptions, and circadian alterations. To mechanistically address this question, we employed both genetic and pharmacological strategies to specifically disrupt the interaction between fibrin and microglia, a key mediator of vascular-induced neuroinflammation.^24^ Genetically, we used *Fgg*^γ390–396A^ knock-in mice, which harbor a targeted mutation in the fibrin(ogen) gamma-chain (Fgg) that prevents binding to the microglial CD11b complement receptors, thus blocking microglial activation by fibrin.^2, 18, 19, 24, 29^ We also used two fibrin-targeting antibodies, 5B8^29^ and THN392, the murine version of the clinical-stage human fibrin-targeting antibody THN391,^27^ which selectively inhibit the fibrin-CD11b interactions with microglia without affecting coagulation.^26, 27, 29^ THN392 exhibit a 100-fold greater affinity for fibrin γ377-395 epitope (Kd = 0.1 nM) than the parent antibody 5B8 (Kd =15 nM).^27^ 5B8 treatment has been shown to reduce microglial activation, loss of cholinergic neurons in the medial septum, as well as inflammatory and oxidative stress responses in the 5xFAD mice.^29^ Here, we show for the first time the beneficial therapeutic effects of *Fgg*^γ390–396A^, 5B8, and THN392 on key features of early AD-related brain network dysfunction, including network hyperexcitability, disrupted sleep-circadian rhythms, and oscillatory EEG slowing.

The affinity matured therapeutic antibody THN391 and its murine variants have been shown to be effective in disease models with prominent fibrin deposition, including multiple sclerosis, fibrin-induced encephalomyelitis, and wet age-related macular edema.^27^ In humans, THN391 was safe and well-tolerated in a completed Phase 1a clinical trial in healthy subjects with no impact on coagulation.^26, 104^ However, the effects of fibrin-targeted therapeutics on early AD-related network dysfunction have not been defined previously. We used the 5xFAD amyloid model which captures the vascular dysfunction and fibrin(ogen) brain deposition^19^ observed in humans with AD^18, 53^ and exhibits prominent AD-related network dysfunction.^105^ These complementary approaches allowed us to mechanistically assess the role of fibrin-induced microglial activation as a previously unrecognized key contributor to AD-related brain network dysfunction.

Consistent with previous findings,^18–20, 53^ we found prominent fibrin(ogen) deposits in the brain parenchyma of AD patients, including accumulation within amyloid plaques, diffuse distribution as perivascular leakage, and presence within some blood vessels (**Figure 1A**). Similar patterns were observed in others amyloid mouse models,^15^ including transgenic 5xFAD and humanized *App*^SAA^ and *App*^NLG^:*APOE4* knock-in mice (**Figures 1B**–**1D**), indicating that fibrin(ogen) deposition is independent of APP overexpression. 5xFAD mice displayed a full array of AD-related brain dysrhythmias (**Figures 2**–**3**) and behavioral abnormalities (**Figure 4**): (1) network hyperexcitability, predominantly during sleep; (2) altered oscillations, characterized by pronounced EEG slowing with increased power in slow-frequency oscillations (delta, theta, and beta) and reduced power in high-frequency oscillations (gamma); (3) marked alterations in sleep architecture, including sleep fragmentation, excessive daytime sleepiness, and nocturnal restlessness; (4) circadian alterations including increased nocturnal hyperactivity; and (5) behavioral abnormalities, such as hyperlocomotion, increased anxiety, and altered spontaneous behavior. Both genetic (*Fgg*^γ390–396A^) and pharmacological antibody-based (5B8 and THN392 antibodies) interventions counteracted these 5xFAD-associated alterations in brain network activity and behavioral alterations, indicating that fibrin-microglia-induced neuroinflammation promotes AD-related brain network and behavioral dysfunction. Mechanistically, fibrin-microglia interactions provide multiple pathways through which neuroinflammation can destabilize synapses,^19^ drive synapse loss,^106^ and cause excitatory-inhibitory imbalance and network hyperexcitability^107, 108^ that undermine learning, memory, and sleep-dependent plasticity^109^. At the molecular level, this is supported by transcriptomic and multi-omic profiling demonstrating that fibrin induces complement (C1q-C3) signaling, oxidative stress pathways (e.g., HMOX1, SOD2), and phagolysosomal and lipid-handling programs (e.g., CTSS, LPL, APOE) in microglia, which together promote synapse elimination, metabolic stress, and impaired neuron-glia coupling.^19, 24, 29^ The companion manuscript by Yan et al. uses calcium imaging to show reduced neuronal hyperactivity by targeting fibrin in the 5xFAD:*Fgg*^γ390–396A^ mice and reports beneficial effects of blocking fibrin-microglia interactions on the restoration of molecular pathways governing microglia-neuron ligand-receptor crosstalk in 5xFAD:*Fgg*^γ390–396A^ mice.^110^ Importantly, the ability of genetic or antibody-mediated disruptions of fibrin-microglia signaling to rescue synaptic^19^ and network dysfunction supports the concept that fibrin-driven synaptic injury^19^ and brain function are reversible and that cognitive decline represents a tractable therapeutic target even after disease onset.

### Vascular dysfunction and network hyperexcitability

Vascular dysfunction, including BBBd and CAA, is increasingly recognized as a key contributor to neural network hyperexcitability in multiple neurological disorders, including AD, epilepsy, and traumatic brain injury.^31–35^ Notably, APP duplications, familial- and CAA-AD linked Aβ mutations (such as the Dutch and Iowa variants), and Down syndrome cause early-onset AD with prominent CAA and higher rates of seizures within familial AD spectrum,^36–40^ suggesting than BBBd and CAA are the pathological drivers of network hyperexcitability. Remarkably, the BBBd subtype is enriched with intracellular Aβ mutations, including the Dutch and Iowa variants, that result in prominent CAA.^16^ However, this subtype is not enriched in PSEN1 carriers, who tend to cluster in subtype 1 (hyperplasticity) and subtype 2 (innate immune).^16^ The high-affinity binding of fibrin(ogen) with these CAA-linked Aβ familial mutations^41, 42^ provides additional mechanistic link between fibrin and network hyperexcitability. Breakdown of the BBB allows blood proteins, such as fibrin(ogen) and albumin, to enter the brain parenchyma, where they can activate microglial and astrocytic inflammatory pathways that promote neuronal excitability. Interestingly, consistent with our results showing that EEG slowing triggers AD-related epileptiform activity (**Figure 3E**), an EEG signature of epileptic activity characterized by a transient slowing of cortical network activity termed paroxysmal slow wave events (PSWEs) has been identified in epileptic patients in cortical regions displaying BBBd and in rodent models with BBBd, including 5xFAD mice.^34^ Notably, infusion of the blood protein albumin triggered PSWEs and increased susceptibility to PTZ-induced seizures,^34, 35^ demonstrating that BBBd and the leakage of blood proteins into the brain parenchyma can lead to hyperexcitability. Albumin has been shown to activate astrocytes through TGF-β signaling, leading to neuronal hyperexcitability, impaired neurogenesis, and epileptiform activity.^32, 33, 111^ In contrast, fibrin(ogen) has not been previously linked before to AD-related network hyperexcitability. We found that AD-related epileptiform activity in 5xFAD mice was reduced by genetic *Fgg*^γ390–396A^ manipulation and fibrin antibody treatments (5B8 and THN392) (**Figures 2**, **5, and 6**), indicating that fibrin-induced neuroinflammation promotes AD-related network hyperexcitability. Furthermore, transcriptomic studies showing that fibrin-dependent microglial activation disrupts microglia-neuron communication networks and engages complement, oxidative and synapse-modifying gene programs that promote circuits toward hyperexcitability.^24, 29, 110^ Together, these findings suggest that both fibrin(ogen) and albumin are not merely markers of BBB leakage in the brain parenchyma but active contributors to AD-related neural hyperexcitability. It is also important to note that BBB leakage–associated blood proteins can actively drive network dysfunction through distinct glial pathways, with albumin acting primarily via astrocytic TGF-β signaling and fibrin acting via microglial CD11b signaling. However, the anti-epileptic effects of blocking albumin or astrocytic TGF-β signaling in AD mouse models have not been directly assess. Altogether, our results confirm the strong mechanistic link between vascular dysfunction and network hyperexcitability and pinpoint fibrin and neuroinflammation as a novel contributor of brain network hyperexcitability.

### Fibrin-induced AD-related brain dysrhythmias and sleep-circadian alterations

AD is characterized by neural hyperactivity detected by MRI, altered oscillatory rhythms, deactivation deficits in the default network, and subclinical epileptiform activity.^30, 76^ These brain dysrhythmias occur in preclinical stages of AD, including in asymptomatic APOE4 and familial AD-mutant carriers, and in individuals with mild cognitive impairment (MCI).^43, 48, 49, 54, 55, 58, 70–75^ For example, MCI patients show early deterioration of gamma oscillations and theta-gamma coupling that progressively deteriorates in AD.^43, 98, 99, 112–114^ Notably, these aberrant patterns of brain network activity seem to directly contribute to AD-related pathogenesis, including increased Aβ^48^ and tau^49^ accumulation, and cognitive decline,^43, 48–51^ suggesting a mechanistic role in AD pathogenesis. AD patients have an increased incidence of seizures and epileptiform activity.^30^ The prevalence of subclinical epileptiform activity in AD patients without seizures is 22–50%^54–57, 115, 116^ and is associated with accelerated cognitive decline.^55^ Mouse models of AD, including APP-Tg^59–68^ and App-KI^60^ mice, also develop network hyperexcitability. AD patients^77, 117, 118^ and related mouse models^98, 99, 112–114^ also show altered brain oscillatory activity, characterized by prominent EEG slowing with increased power of slow-frequency oscillations and reduced power of high-frequency oscillations,^77, 117, 118^ which has been also associated with cognitive decline.^119^ As in humans,^30, 43^ we found prominent EEG slowing (increased power in slow-frequency oscillations and decreased power in high-frequency oscillations) in 5xFAD mice that was ameliorated by *Fgg*^γ390–396A^ (**Figure 2E**) and chronic THN392 treatment (**Figures 6F-6H**), indicating that fibrin-induced neuroinflammation promotes AD-related EEG slowing.

We also explored the aperiodic or non-oscillatory component of the power spectral density, which has been recently linked to brain function.^77, 79–85^ Notably, we found higher aperiodic exponents or decays, which correspond with resting brain states like sleep^82^ or anesthesia,^81^ in 5xFAD mice, which were also ameliorated by *Fgg*^γ390–396A^ (**Figure 2H**) and chronic THN392 treatment (**Figure 6D**). Notably, AD patients appear to show either no change^77^ or a reduction in the aperiodic decay^120^, which has been interpreted as reflecting reduced inhibitory tone, since anesthesia induced by the GABAA agonist propofol increases the aperiodic decay^81^. Our interpretation differs, as we observed increased epileptic activity during resting states (**Figure 2C**) characterized by higher aperiodic decay (**Figure 2H**), including quiet wakefulness and sleep. This view is consistent with the notion that resting brain states are strongly associated with epileptic activity in AD and other neurological disorders.^30^ Although these resting states may involve enhanced inhibitory tone, they are also marked by large-scale synchronized slow oscillations, which can create conditions that favor pathological hypersynchrony in AD.^30^ Thus, our aperiodic results are consistent with the established notion of EEG slowing and network hyperexcitability in AD, and we propose that higher aperiodic decay, rather than indicating increased inhibitory tone, should be interpreted as a shift toward resting-state brain activity patterns.^82^

Sleep^44^ and circadian rhythm^45^ disruptions are common and early features of AD, often preceding cognitive decline.^44^ AD patients frequently experience fragmented sleep, increased nighttime wakefulness (nocturnal restlessness), daytime sleepiness, and altered timing of sleep-wake cycles.^46, 121^ These disturbances have been linked to accelerated amyloid deposition and neurodegeneration, impaired glymphatic clearance of toxic proteins, and neuroinflammation,^46^ suggesting that sleep and circadian dysfunction are not only symptoms or comorbidities but also key contributors to AD pathogenesis. A recent study found that insomnia increased the risk of cognitive decline in MCI and dementia patients, as well as anxiety, greater white matter hyperintensities, and amyloid-PET burden.^86^ Additionally, insomnia has also been associated with cardiovascular and cerebrovascular disease.^122–126^ We observed similar marked alterations in sleep architecture in 5xFAD mice, including sleep fragmentation, excessive daytime sleepiness, arousal instability, and nocturnal restlessness, which were also ameliorated by *Fgg*^γ390–396A^ (**Figures 3A and 3B**). Interestingly, altered oscillatory activity, network hyperexcitability, and sleep seem to be pathologically linked in AD. For example, sleep stages, including REM and NREM, have been closely associated with increased epileptic activity in AD patients^55, 58, 91, 92^ and related animal models^127^, likely due to the elevated slow-frequency oscillations which increases neuronal synchrony and promotes the emergence of subclinical epileptiform discharges. NREM-associated hyperexcitability and reduced NREM bout duration may exacerbate cognitive decline by disrupting memory consolidation and sleep-dependent neural homeostasis.^128^ Altogether, the tight interdependence among brain network dysrhythmias—linking oscillatory activity with epileptic discharges and sleep stages with both oscillations and spikes—suggests a shared underlying mechanism. Our findings indicate that fibrin-induced neuroinflammation may serve as a central driver of these abnormalities. Furthermore, these insights suggest that sleep architecture should be explored in clinical trials, as it provides an accessible biomarker that correlates with a diverse array of AD-related brain dysrhythmias.

### Fibrin-induced neuroinflammation as a therapeutic target for AD

Fibrin-induced neuroinflammation has emerged as a promising and novel therapeutic target in AD due to its critical role in driving neurovascular dysfunction and neuroinflammation.^1^ Fibrin deposits activate microglia through CD11b complement and integrin signaling, triggering chronic neuroinflammation and contributing to synaptic dysfunction and behavioral alterations.^19, 24, 25, 129^ Here, we show that this pathway plays a central role on AD-related brain hyperexcitability, altered oscillations, sleep, and circadian changes. This is supported by previous studies demonstrating that fibrin-CD11b signaling drive microglia-mediated spine elimination and cognitive impairment^19, 24^ and positions fibrin-induced neuroimmune remodeling upstream of the network and sleep phenotypes observed here. Thus, targeting fibrin or its pathological interactions with microglia via immunotherapy against fibrin or other strategies to preserve BBB integrity represents a promising disease-modifying approach for reversing inflammatory and vascular cascades that contribute to AD, particularly in AD patients with BBBd. Importantly, the development of fibrin-specific monoclonal antibodies that selectively target pathological fibrin without affecting systemic coagulation offers a promising translational tool to advance fibrin-targeting therapies to the clinic.^1, 26, 104^ Indeed, TNH391 has recently shown a favorable safety profile in a Phase 1a clinical trial involving healthy volunteers with a prolonged half-life of approximately 40 days and no impact on coagulation. Phase 1b clinical trials are ongoing to evaluate THN391 in patients with AD and diabetic macular edema. Our data support the use of neural network functional outcomes for evaluating the efficacy of fibrin-based antibody therapies.

We would like to highlight several limitations of this study. First, our therapeutic intervention was restricted to symptomatic 5xFAD mice. While we observed significant beneficial effects, it remains unclear whether a preventive strategy initiated at the onset of BBBd and fibrin deposition would be equally or even more effective, as this therapeutic approach may require significant BBBd to achieve efficacy. Similarly, while the 5xFAD model is characterized by robust BBB leakage and amyloidosis, future studies should investigate whether models featuring both amyloid and tau pathology—but lacking overt BBBd —would derive similar benefits. Regarding our functional outcomes, we focused on spontaneous behavior and brain network functions rather than direct assessments of spatial memory. Although previous work has shown that *Fgg*^γ390–396A^ improves spatial memory in 5xFAD mice,^19^ the long-term impact of anti-fibrin antibody therapy on learning and memory remains to be determined. Given the improvements we observed in brain function and spontaneous activity, we anticipate that these benefits would translate into cognitive benefits. Finally, while we confirmed that both sexes in 5xFAD mice exhibit similar deficits and responses to *Fgg*^γ390–396A^, our anti-fibrin antibody cohorts were not specifically powered to detect sex-dependent differences in treatment efficacy.

## ACKNOWLEDGEMENTS

We thank Allan Villanueva, Rhodora Gacayan, and Eilidh MacDonald for technical assistance. The study was in part supported by a sponsored research agreement from Therini Bio to the Gladstone Institutes (K.A. and J.J.P.), Warren Alpert Distinguished Scholar Award (Z.Y.), Brightfocus Postdoc Fellowship Award A2021019F (Z.Y.); Kaganov Scholarship for Excellence in Neuroscience (Z.Y.); philanthropic gifts from Edward and Pearl Fein, Robert Hamwee, the Dolby Family Fund, the Spangler Foundation, the Simon Family Trust (K.A.); and the US National Institutes of Health (NIH) grants RF1AG062234, P01AG062629, R01AG073082, and AG082147 (J.J.P.), K01-AG083732 (S.R.M.), K99 NS126707 (A.S.M), R35 NS097976 and R35 NS143067 (K.A.), and RF1AG06926 (K.A., J.J.P., and MH.E.).

## AUTHOR CONTRIBUTIONS

Conceptualization: K.L., Z.Y., A.S.M., Y.Z., D.M., J. Stavenhagen, A.B.K., M.H.E, K.A., and J.J.P; Data curation: K.L. and J.J.P.; Formal analysis: K.L., D.M., E.B., and J.J.P.; Investigation, mouse generation: K.L., Z.Y., A.S.M., R.M.A., B.C., M.D.P.S.A., Y.Z., J. Shin, E.B., K.S., J.H., J.K.R., K.A., and J.J.P; Immunohistochemistry: K.L., Y.Z., J. Shin, J.S.R., I.C., and J.J.P.; *In vivo* pharmacology: K.L., S.R.M., P.N., Z.Y., A.S.M., R.M.A., B.C., M.D.P.S.A., J.K.R., J. Stavenhagen, A.B.K., K.A., and J.J.P; EEG recordings: K.L. and J.J.P.; Machine learning behavior: K.L., S.R.M., D.M., P.N., N.K., and J.J.P; Software/coding: K.L., S.R.M., E.B., D.M., P.N., F.J., K.M., N.K., A.A., R.S., and J.J.P; Data analyses: K.L., S.R.M., Y.Z., E.B., D.M., P.N., N.K., and J.J.P; Data interpretation: K.L., S.R.M., P.N., Z.Y., A.S.M., J. Stavenhagen, A.B.K., I.C., M.H.E, K.A., and J.J.P; Resources: J. Stavenhagen, A.B.K., K.A., J.J.P.; Visualization and statistics: K.L. and J.J.P.; Writing, original manuscript: K.L. and J.J.P.; Writing, review & editing: all authors; Supervision: K.A. and J.J.P.; and funding acquisition: K.A., M.H.E., and J.J.P.

## STAR★METHODS

## KEY RESOURCES TABLE

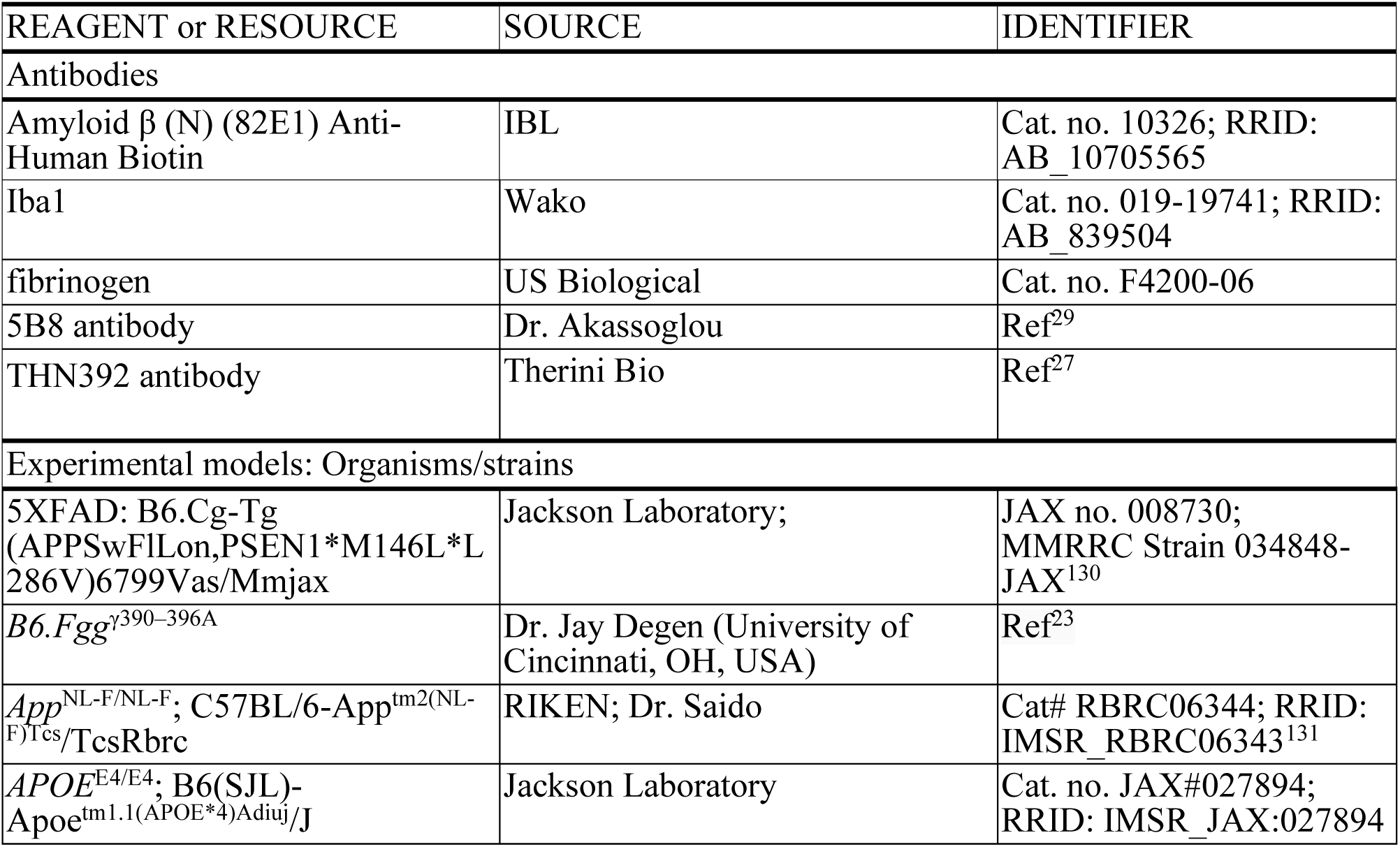
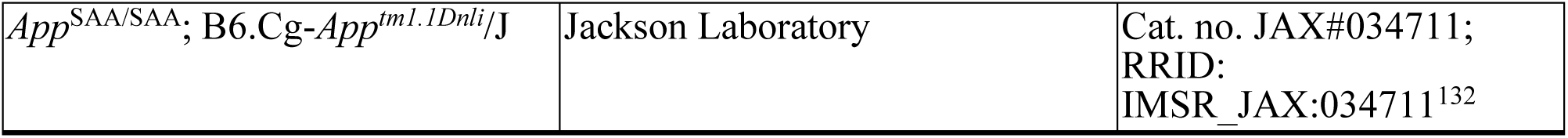

## EXPERIMENTAL MODEL AND SUBJECT DETAILS

### Mice

5xFAD mice were from The Jackson Laboratory (Jax 0087030).^130^ We studied male and female heterozygous 5xFAD mice overexpressing both human amyloid beta precursor protein (APP) with the Swedish (K670N, M671L), Florida (I716V), and London (V717I) FAD mutations and human presenilin 1 (PS1) with the M146L and L286V FAD mutations on the C57BL/6J background.^130^ We used male 5xFAD mice for breeding since maternal Thy1 promoter expression results in reduced transgene expression and lower amyloid.^133^ *Fgg*^γ390–396A^ mice were from Dr. Jay Degen (University of Cincinnati, OH, USA) and bred in the Akassoglou Lab.^23^ Heterozygous 5xFAD mice were crossed with wildtype (WT) mice to produce heterozygous 5xFAD and WT mice on the C57BL/6J background. Homozygous *Fgg*^γ390–396A/γ390–396A^ mice were crossed with 5xFAD:*Fgg*^γ390–396A/γ390–396A^ mice to produce 5xFAD:*Fgg*^γ390–396A/γ390–396A^ and *Fgg*^γ390–396A/γ390–396A^ mice. Homozygous *Fgg*^γ390–396A/γ390–396A^ mice are referred to as *Fgg*^γ390–396A^ mice. Experimental mice of the same sex were group-housed with access to water and food *ad libitum* in a controlled environment with a 12-hour light-dark cycle.

*App*^NL-F^ mice were obtained from Drs. Takaomi Saido and Takashi Saito (RIKEN Brain Science Institute, Japan).^131^ *APOE4* mice were obtained from The Jackson Laboratory (Jax 027894). We crossed *App*^NL-F/WT^:*APOE^E4/WT^* to produced *App*^NL-F/NL-F^:*APOE^E4/E4^* mice. We studied male and female homozygous *App*^NL-F/NL-F^:*APOE^E4/E4^* mice expressing humanized Aβ and the familial Alzheimer’s disease (FAD) Swedish (K670N, M671L) and Iberian (I716F) mutations and human *APOE^E4/E4^* on the C57BL/6J background. Homozygous *App*^NL-F/NL-F^:*APOE^E4/E4^* mice are referred to as *App*^NL-F^:*APOE4* mice. *App*^SAA^ mice were obtained from The Jackson Laboratory (Jax 034711). We studied male and female homozygous *App*^SAA/SAA^ mice expressing humanized Aβ and the familial Alzheimer’s disease (FAD) Swedish (K670N, M671L), Artic (E693G), and Austrian (T714I) mutations on the C57BL/6J background.^132^ Homozygous *App*^SAA/SAA^ mice are referred to as *App*^SAA^ mice.

EEG and behavioral experiments were performed in sex-balanced groups by investigators who were unaware of the genotype or treatment groups of the mice. All mice were bred and housed at the Gladstone Institutes vivarium, a UCSF AAALAC-approved facility. All mouse experiments were approved by the Institutional Animal Care and Use Committee of the University of California, San Francisco, and were conducted in accordance with the NIH guidelines for the care and use of laboratory animals.

### Human tissue

Brain tissue from AD patients and age-matched control donors was obtained from the UCLA Department of Pathology. AD neuropathology was evaluated by a neuropathologist using the ABC score (National Institute on Aging and Alzheimer’s Association Research Framework criteria) (MPID22265587), cases used were all A3, B3, C3. Relevant information such as age, sex, ethnicity, brain weight, and postmortem interval (PMI) were recorded when available. No cases with imaging or gross findings consistent with large vessel territorial infarction, hemorrhage, primary or metastatic neoplasms, or CNS infection were included. Cases with histological evidence of hypoxic-ischemic brain injury were excluded. Tissue blocks underwent immunohistochemical assessment, including H&E and Nissl stains to confirm tissue integrity and the absence of microinfarcts or other focal pathologies.

## METHOD DETAILS

### 5B8 and THN392 antibody treatments

5B8 and an isotype-matched IgG2b control (MPC-11, BioXCell) antibodies were dosed i.p. every other day (48 hours) for 2 weeks (8 total doses) at 30 mg/kg. THN392 and an isotype-matched IgG2b control (MPC-11, BioXCell) antibodies were dosed biweekly (three and four days apart) for 13 weeks (25 total doses) at 30 mg/kg. 5B8 was produced as previously described^29^. THN392, the murine IgG2b version of the humanized therapeutic antibody THN391, was produced in CHO-HP cells by GenScript (Piscataway, NJ) for Therini Bio.^27^ It was purified by MabSelect^TM^ PrismA Protien A chromatography and is 98% pure by SEC-HPLC. Endotoxin is ≤0.1 EU/mg.

### EEG transmitter implantation and telemetry recordings

Telemetric EEG/EMG transmitters (HD-X02) (Data Sciences International) were implanted according to the manufacturer’s surgical protocol for EEG and neck electromyography (EMG) in mice. In brief, mice were anesthetized with 3% isoflurane oxygen which was reduced to 0.5-1% during surgery; local anesthetized with lidocaine, for analgesia, buprenorphine (0.05 mg/kg) was given before surgery and ketoprofen (5 mg/kg) was given at the end of the surgery. A 2–3-cm incision was made from posterior of the eyes to the scapulae, and the transmitter was implanted into a subcutaneous pocket along the dorsal flank. Two biopotential EMG leads were placed in the cervical trapezius muscles in the dorsal region of the neck. EEG leads were placed in the left and right posterior parietal cortex at -2 AP, and (+/-2) ML.

Continuous EEG, EMG, activity, and temperature signals were recorded within a normal mouse vivarium housing room 24 hours a day for several weeks using the Data Science International hardware and Ponemah software. Mice were placed into new clean cages the afternoon before the start of the first day (starting at 7 am) of the recording to reduce elevated activity associated with a novel environment, exploration, and nesting within the first 12 hours of being placed in a new cage. Mice were monitored daily. Recordings were analyzed according to ZT time based on the 12 hour light/dark schedule (starting at 7 am). EEG, EMG, and activity signals were collected at 500 Hz and temperature was collected at 10 Hz. Following completion of the experiment, recordings were exported as edfs. and analyzed for brain wave oscillations, sleep time and transitions, locomotor activity, epileptic spikes, and temperature using Spike2 software (Cambridge Electronic Design Limited), excel, SPSS, and GraphPad Prism.

Large noise events were filtered out of the EEG, EMG, activity, and temperature data if it passed a particular threshold (well above normal physiological signals) for EEG/EMG a threshold of >2 mV or <-2 mV, activity (<0 counts) temperature (≤25 degrees) was used and noise removed using a Spike2 script (Unclip) and later a custom made python script. When recordings from the same mouse needed to be merged such as in the instance of days separated by a stop in the recording for a treatment then either the custom python script was used and/or a Spike2 script (Merge Files (before 08/24 update)).

Oscillatory power bands were analyzed in Spike2 using the spectral band power analysis function and/or a custom made Spike2 script which calculated and created all the oscillatory frequency power bands at once in separate virtual channels. As the EEG recording were all collected at 500 Hz the frequency resolution for all the frequency bands was set to 0.5 Hz for an FFT of 1024 points. Following frequency band analysis, a FIR digital filter, high pass filter, was applied to the EEG channel with the high pass set at 6.3 and the transition gap set at 5.0, and was consistent across all experiments. Spike2 was used for spike analysis using the spike template WaveMark search. Spikes were determined to be abnormal activity well beyond the normal EEG activity, (10x the SD of the baseline taken at three, 2-minute intervals in the beginning middle and end of the recording with little to no spikes.) and 5 to 80 ms in length and having a shape and frequency power analysis consistent with epileptiform spiking and opposed to noise or artifacts. Data was exported at 500 Hz then averaged over 1-minute bins that were then averaged into 15-minute bins over 7 days (1 week) 24 hours per day based on ZT time (0 to 24 hours, 7am to 7am) for statistical analysis. Spike2, Excel, SPSS and Prism were utilized for the analysis and graphs. Binning was applied to parameters such as locomotor activity (Rest-to-active level; Figures 2E, 2H, 6E–6I, S1) and oscillatory power (Figures 3E, 5F, 6J–6N, S4A) based on the distribution of observations. For example, Rest-to-active levels were divided into five equal-frequency bins, with each bin containing 20% of the data: bin 1 representing the lowest locomotor activity and bin 5 the highest. Statistical analysis generally consisted of generalized linear mixed model accounting for repeated measures, individual differences (random factor), and fixed factors such as Day/Night, activity bin, etc. dependent on the experiment with Bonferroni posthoc tests to account for multiple comparisons.

One hour of the EEG recordings was taken from the light and dark phase to be used for analyzing the aperiodic spectral features (day 2; 10 AM and 10 PM). Across 10 second bins, mean spectra were generated using the function mtspecgramc from the Chronux toolbox^134^ (5 tapers; 0.5 second non-overlapping window). A frequency range of 4-70 Hz was used for periodic analysis while 15-70 Hz was used to analyze the aperiodic exponent. Aperiodic and periodic components of the average spectra were parameterized using the FOOOF algorithm.^84^ Settings for the algorithm were as follows: peak width limits = [3 30]; max number of peaks = [inf], min peak height = [0]; peak threshold = [1]; aperiodic mode = ‘fixed’. If a periodic peak was identified in the following frequency ranges the peak log-transformed power was taken as well as the area under the curve (AUC) between the linear periodic signal and aperiodic exponent: theta = 6-12 Hz; beta low = 11-15 Hz; beta high = 15-25 Hz; low gamma = 25-40 Hz; high gamma = 40-70 Hz.

Sleep analysis was performed in Spike2 using a script (RatSleepAuto) at the 5 second epoch with the default settings for sleep detection. REM detection criteria was defined as alpha = 2.5, beta = 4 (REM criteria: t:d > 1.#IO; Mn.r.emg <0.011) while NREM detection criterion was defined as delta = 3, epsilon = 3 with EMG faded gamma = 3, Mu = 1, faded omega = 1.5; (EEG >0.035; Delta > 0.022; emg < 0.007). Transitions into the sleep states and time in the sleep states were calculated based on the 5 second epoch data for the averaged 7 day means for each 15-minute bin over 24 hours. The specific transitions, bout length, number of bouts, epileptiform spikes per sleep stage, and specific bout length counts were analyzed from the means for each of the 7 days.

### Open Field Test

A one-hour novel open field (OF) was conducted on forty-seven mice, age range 7-10 months (Ave = 8.5 months) with four groups and similar female:male ratios in the 5xFAD (n = 13; 7:6), 5xFAD:Fgg^γ390–396A^ (n = 12; 6:6), Controls (n = 22; 10:12) mice (**Figure 4**) and THN392 behavior cohort; 7-8 month-old mice TNH292 (n = 5) or IgG2b (n = 5) antibody (**Figure 6**). Circular OF chambers were used with a 12” inner diameter. Male or female mice were tested in alternating blocks to ensure balanced experimental groups throughout the day. The chambers were cleaned with 70% ethanol between trials. All OFs were conducted during the light cycle of the mice in a separate behavioral room. Mice were allowed to habituate for at least 1 hour before running the OF. To optimize exploratory activity lower lux levels in the range of 60-70 lux were used for illuminating the chambers. Mice were video recorded from below using EthoVision video tracking system (Noldus Information Technology). Distance moved within the arena, percentage of time moving or stationary, and zone time (wall or center) were measured using EthoVision software. Statistical analysis was conducted in GraphPad Prism and consisted of two-way ANOVAs with Tukey’s test to correct for multiple comparisons.

### Machine learning (ML) VAME analyses of spontaneous behavior

We used our recently developed ML-VAME^25, 103^ approach for assessing spontaneous behavior in an open arena. Briefly, as recently described^25^, we will collect 60-min videos for each mouse after 8 weeks of THN392 or vehicle treatment. We used DeepLabCut (DLC)^135, 136^ for pose estimation of 11 mouse body parts (virtual skeleton) and VAME^25, 103^ for behavioral segmentation into 30 ethologically-relevant behavioral motifs.

### Machine Learning Spatial Analysis

Our ML VAME analysis was expanded to extract spatial data, center vs periphery, for each of the motifs and communities for the one hour OF. Zone selection: the arena center coordinates and radius were extracted using the Hough Gradient Method (OpenCV) applied to the first frame of each video. Pose estimates from DeepLabCut, based on the belly point, were used to assign each frame to spatial zones according to distance from the center of the chamber — The center zone was defined as ≤0.6 of the arena radius, and the periphery as >0.6-1.0 of the radius.

To quantify behavioral changes over time, we computed the percentage of time spent in the arena center for each motif or community, in each genotype group. The percent time in the arena center was computed for each motif/community by dividing the number of motif/community-labeled frames occurring in the central zone by the total number of frames for that motif/community, yielding a per-motif percent time-in-center measure. These values were calculated at both the session level and at 10-minute time bins for temporal comparisons. Outlier sessions were excluded if they were pre-identified as invalid and only one outlier was excluded from the data set.

Probability density heatmaps: to assess spatial patterns of behavioral motif/community usage, we computed two-dimensional probability density maps. For each session, zone coordinates were extracted from DeepLabCut-derived positional coordinates and aligned with the corresponding motif/community labels from VAME. When specific time windows subsets of the data were specified, position and vectors were restricted to the selected frame range. For each defined state of motif or community sets, the concatenated coordinates were used to estimate a continuous probability distribution via Gaussian kernel density estimation (KDE; SciPy, Scott’s bandwidth method). Densities were evaluated on a grid of 200 × 200 bins covering the spatial extent of the arena. To facilitate visualization, density values were normalized by the maximum probability within each state and masked at <10% of the maximum. The resulting density maps were plotted as contour heatmaps using the inferno colormap (Matplotlib), with probability scaling consistent across groups when comparing genotypes. Colorbars indicate the relative usage probability, and axes correspond to arena coordinates. The group-level KDEs were computed and visualized separately to highlight genotype-specific motif and community usage distributions. Statistical analysis for the OF spatial analysis consisted of two-way ANOVAs with post-hoc Tukey’s multiple comparisons test for the motif and community analysis.

### Histology and immunohistochemistry

Mice were anesthetized and transcardially flush-perfused with 0.1 M phosphate buffer (PB), and brains were extracted and drop-fixed in 4% phosphate-buffered paraformaldehyde at 4 °C for 24 hours. After rinsing with PB saline (PBS), brains were stored in PBS. For sectioning, brains were transferred to 30% sucrose in PBS at 4 °C for 18–24 hours before being frozen and coronally sectioned with a sliding microtome. Ten subseries of free-floating sections (30 μm) were collected per mouse and kept at –20 °C in cryoprotectant medium until use. Each subseries contained sections throughout the rostrocaudal extent of the forebrain.

Brain sections were washed 3 times for 10 min with PBS to remove cryoprotectant medium and once with 0.5% Triton X-100 (PBTx) to permeabilize the tissue. Endogenous peroxidases were blocked with a 15-min incubation with 3% hydrogen peroxide and 10% methanol in PBS. Sections were subsequently washed three times for 10 min each in PBSTx. Nonspecific binding was blocked with a blocking solution containing 10% normal donkey serum (Jackson ImmunoResearch, 017-000-121) and 0.2% gelatin (Sigma-Aldrich, G2500) in PBSTx for 1 hour at room temperature. Brain sections were incubated overnight with rabbit anti-Iba1 (Wako 019-19741, 1:500), mouse anti-human amyloid beta (N) (82E1) Biotin (IBL 10326-S, 1:500), sheep anti-human fibrinogen (US Biological F4200-06, 1:300). After three washes in PBST, sections were incubated in secondary antibodies for 1 hour at room temperature, which include donkey anti-rabbit Alexa488 (Invitrogen A-21206, 1:500), streptavidin Alexa350 (Invitrogen S-11249, 1:500), donkey anti-sheep Cy3 (Jackson ImmunoResearch 713-165-003, 1:500). Sections were washed three times in PBST before mounting and cover slipping with ProLong Diamond Antifade Mountant (Invitrogen P36965). All immunofluorescent images are obtained from a fluorescence microscope (Keyence BZX810) at 40x magnification.

## QUANTIFICATION AND STATISTICAL ANALYSIS

Statistical analyses and graphs were done with SPSS (28.0.1.1), MATLAB (2018b), or Prism 10. Statistical tests and *p* values are indicated in the figure legends. Briefly, Generalized Linear Mixed Model (GLMM), accounting for repeated measures (15 min timepoints for 24 hours averaged over seven days for the circadian cycle analyses; 10 s timepoints for 1 hour during day or night for the aperiodic analyses), fixed factors (day/night, locomotion levels, power levels), and random factors (mouse), along with a Bonferroni post hoc test for multiple comparisons, was used to assess EEG outcomes. Repeated two-way ANOVA was used to assess genotype in behavioral outcomes. *N* number of mice are indicated in the figures and figure legends. Values are mean ± SEM. Null hypotheses were rejected by double-tailed tests with an alpha value of 0.05.

## SUPPLEMENTAL FIGURES

**Figure S1.**
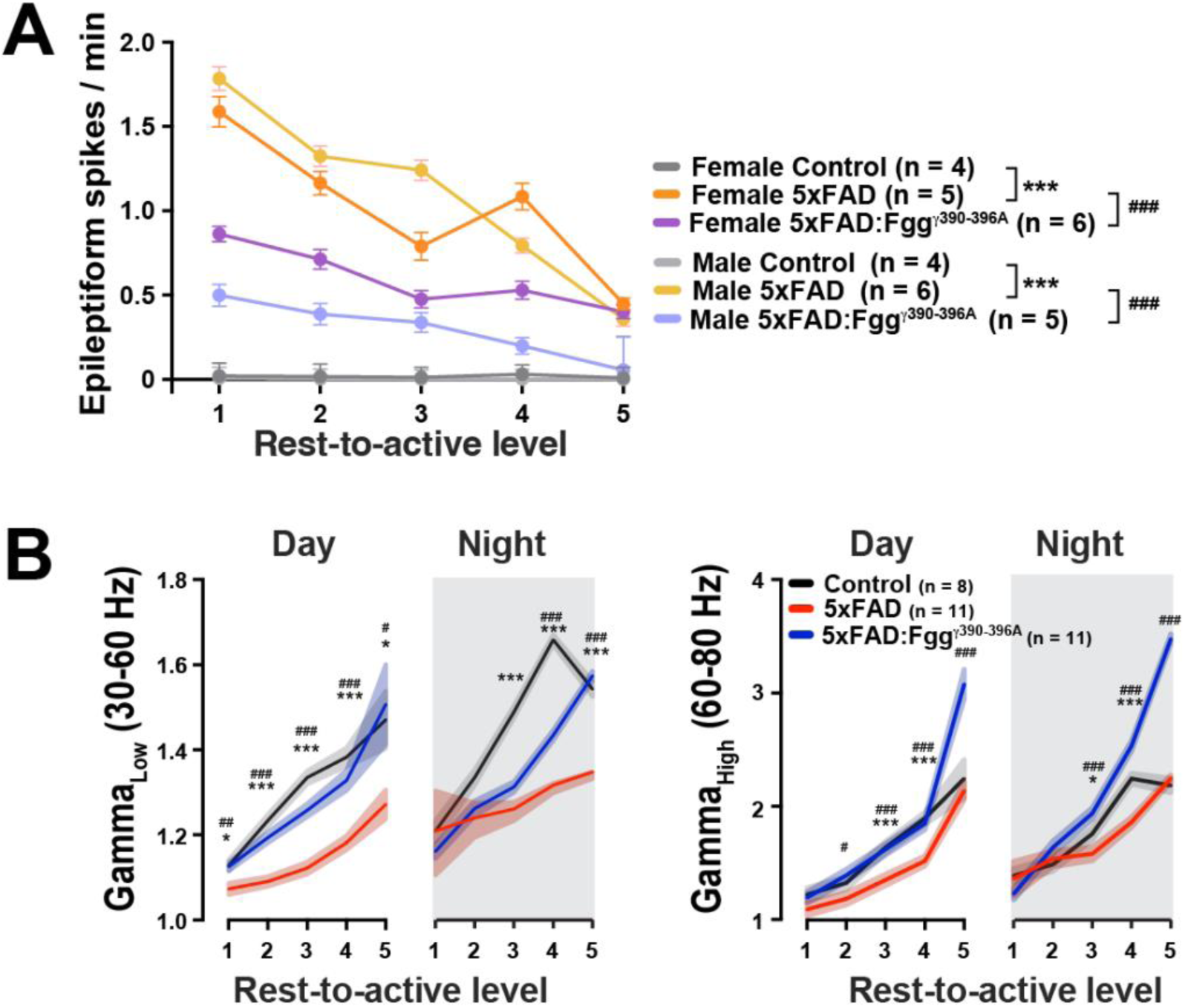
Related to Figure 2. 5xFAD-dependent alterations in both low and high gamma brain oscillations are ameliorated by blocking fibrin–microglia interactions in 5xFAD:*Fgg*^γ390–396A^ mice. 10–13-month-old sex-mixed control (n = 8), 5xFAD (n = 11), and 5xFAD:*Fgg*^γ390–396A^ (n = 11) mice were implanted with a wireless EEG/EMG transmitter to continuously monitor cortical brain activity for 7 days. (**A**) Epileptiform spikes by locomotor activity level (rest-to-active) and sex. Groups included: Control (n = 8; 4 females, 4 males), 5xFAD (n = 11; 5 females, 6 males), and 5xFAD:*Fgg*^γ390–396A^ (n = 11; 6 females, 4 males) mice. Epileptic spikes were strongly modulated by locomotion in 5xFAD and 5xFAD:*Fgg*^γ390–396A^ mice, with periods of low locomotor activity having higher epileptic spike rates and vice versa. Relative to 5xFAD mice, 5xFAD:*Fgg*^γ390–396A^ mice had reduced epileptic activity. (**B**) Low gamma (30-60 Hz) and high gamma (60-80 Hz) power are both reduced during the day and night in 5xFAD mice compared to controls, but rescued in the 5xFAD:*Fgg*^γ390–396A^ mice. Values are mean ± SEM; P values by Generalized Linear Mixed Model (GLMM) accounting for repeated measures, individual differences (random factor), fixed factors (day/night and activity levels), and Bonferroni post hoc test for multiple comparisons. \**p* < 0.05, \*\**p* < 0.01, and \*\*\**p* < 0.001 for Controls vs. 5xFAD; ^#^*p* < 0.05, ^##^*p* < 0.01, and ^###^*p* < 0.001 for 5xFAD vs. 5xFAD:*Fgg*^γ390–396A^.

**Figure S2.**
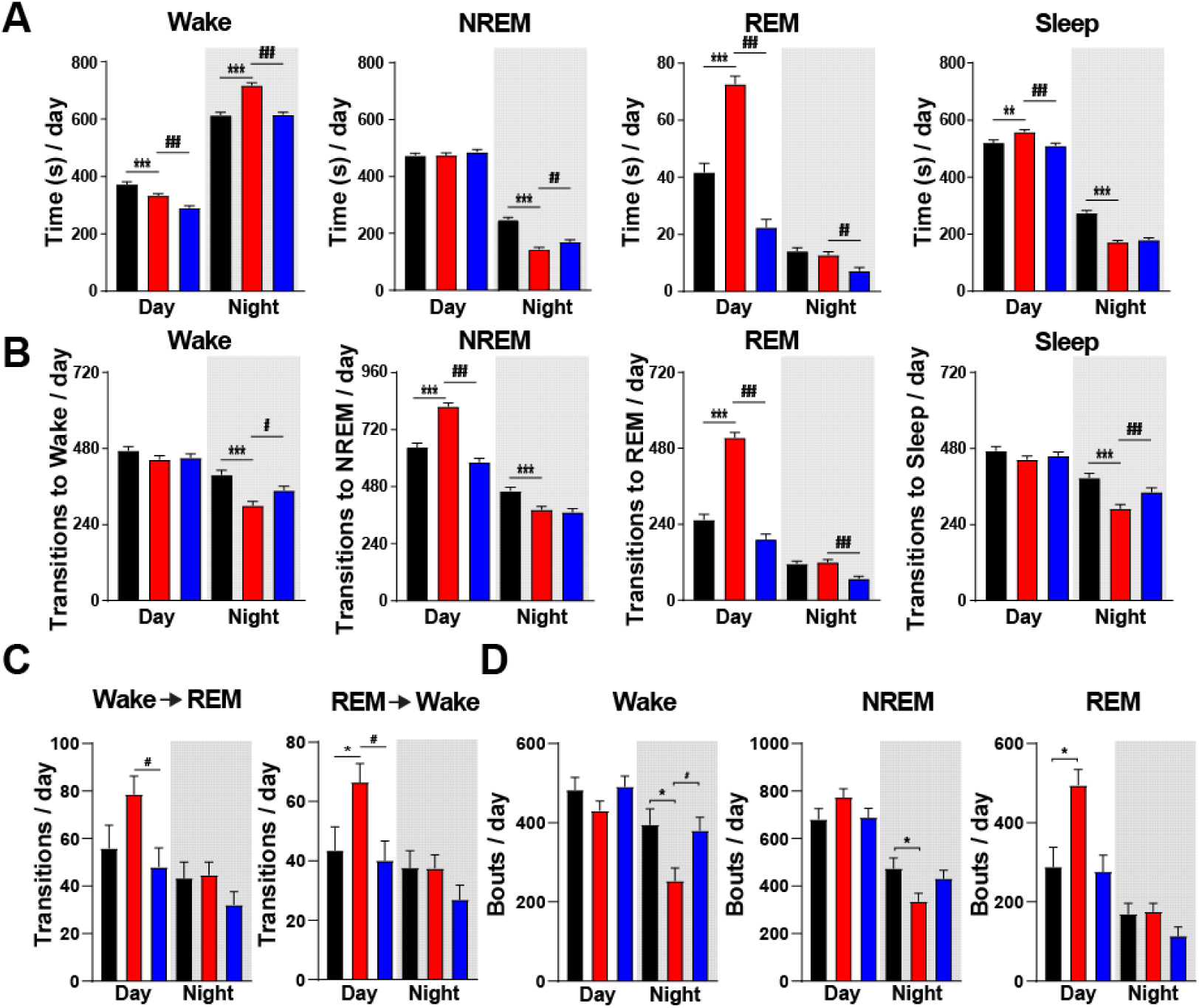
Related to Figure 3. 5xFAD-dependent sleep alterations are rescued by blocking fibrin–microglia interactions in 5xFAD:*Fgg*^γ390–396A^ mice. (**A**) Total circadian time per average 7-day 15 min bin in Wake, NREM, REM, and Sleep (NREM & REM). (**B**) Total average transitions to Wake, NREM, REM, and Sleep per seven-day average. (**C**) Specific transitions from Wake to REM, and REM to Wake during day and night. (**D**) Average total number of bouts in Wake, NREM, and REM in the day and night. Values are mean ± SEM; P values by Generalized Linear Mixed Model (GLMM) accounting for repeated measures, individual differences (random factor), fixed factors (day/night and activity levels), and Bonferroni post hoc test for multiple comparisons. \**p* < 0.05, \*\**p* < 0.01, and \*\*\**p* < 0.001 for Controls vs. 5xFAD; ^#^*p* < 0.05, ^##^*p* < 0.01, and ^###^*p* < 0.001 for 5xFAD vs. 5xFAD:*Fgg*^γ390–396A^.

**Figure S3.**
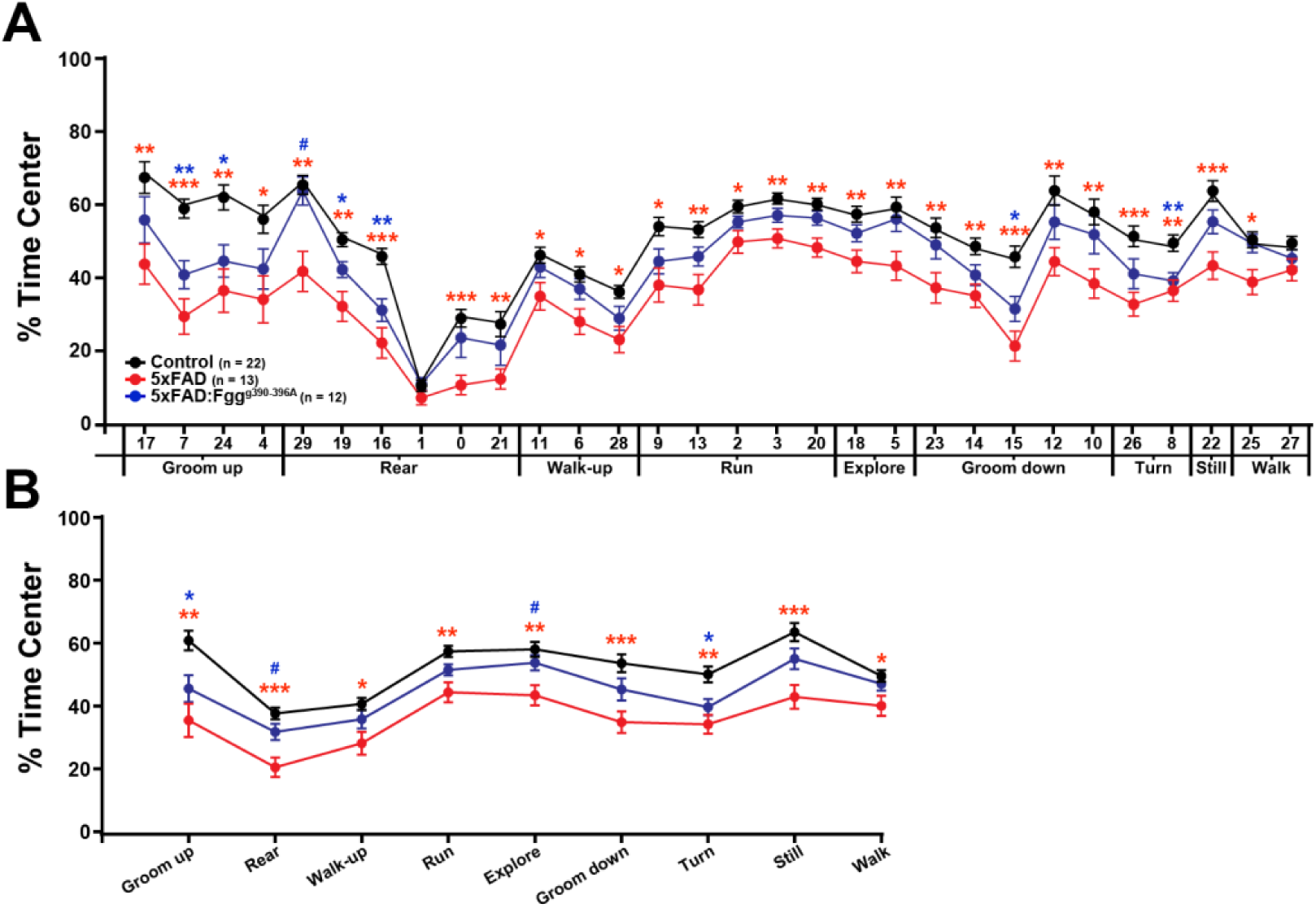
Related to Figure 4. Overall open field spatial analysis shows significantly altered behavior in the center of the arena in 5xFAD mice and recovery across behavioral motifs and communities by blocking fibrin-microglia interactions. (**A-B**) The overall % of time in the center of the open field for each behavioral motif (A) and across communities (B) in control, 5xFAD, and 5xFAD:*Fgg*^γ390–396A^ mice. Values are mean ± SEM; P values by repeated two-way ANOVA with Tukey’s test for multiple comparisons (A-B); \**p* < 0.05, \*\**p* < 0.01, and \*\*\**p* < 0.001 (red asterisk) for Controls vs. 5xFAD, (blue asterisk) for Control vs. 5xFAD:*Fgg*^γ390–396A^; ^#^*p* < 0.05, ^##^*p* < 0.01, and ^###^*p* < 0.001 for 5xFAD vs. 5xFAD:*Fgg*^γ390–396A^.

**Figure S4.**
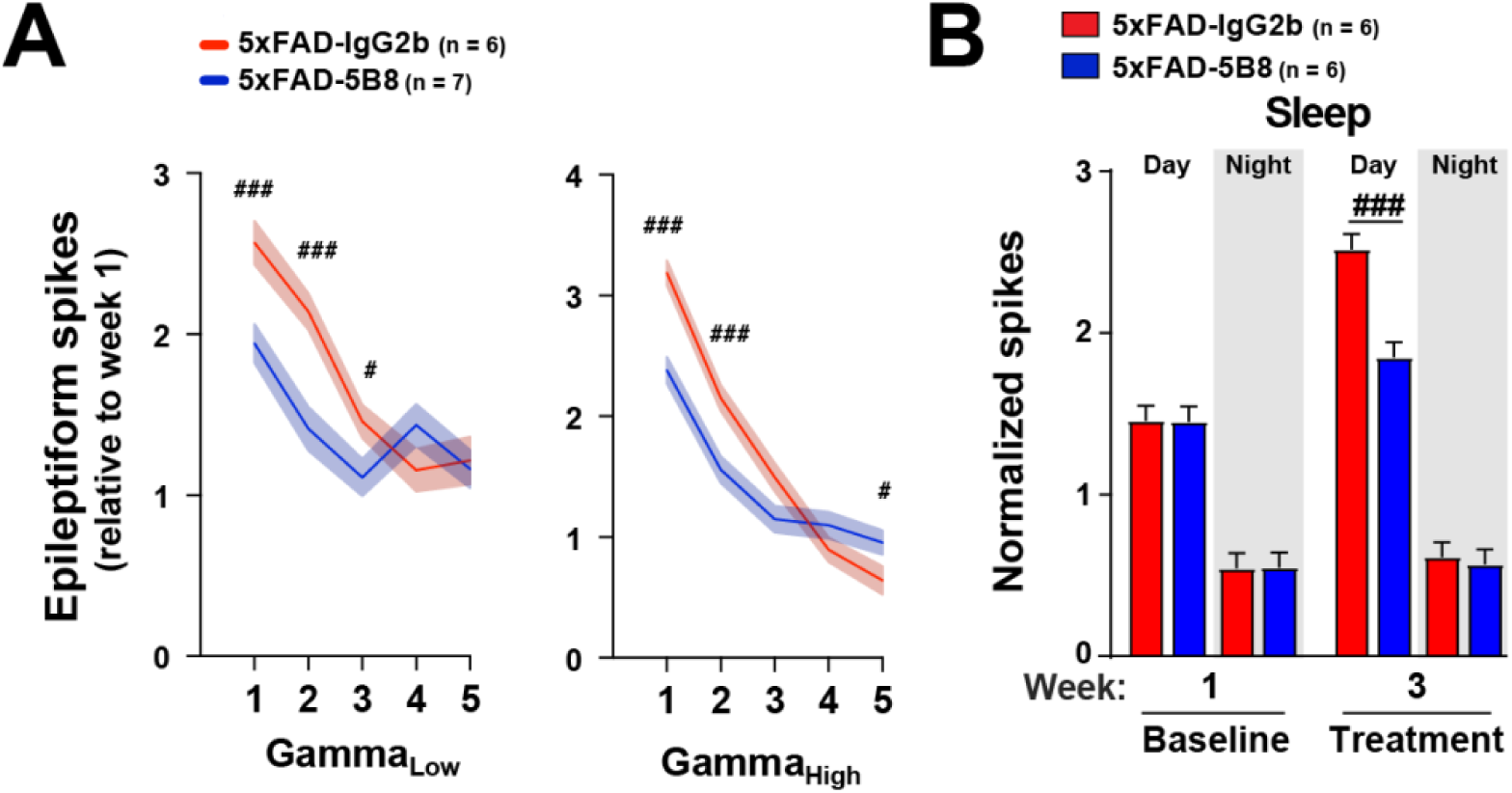
Related to Figure 5. Treatment with the anti-fibrin antibody 5B8 mitigates alterations in Epileptiform spikes observed in low and high gamma, and 5xFAD sleep-dependent alterations. (A) Epileptic activity per oscillatory power level (low-to-high). 5B8 treatment decreased epileptic activity particularly during periods with low power of both low (30-60 Hz) and high (60-80 Hz) fast-frequency gamma oscillations in 5xFAD mice. (B) Epileptic activity during Sleep in 5B8 treated 5xFAD mice. 5B8 treatment decreased epileptic activity during the day in which the majority of epileptiform spikes occur. Values are mean ± SEM; P values by Univariate General Linear Model (UGLM) accounting for repeated measures, fixed factors (power levels for A; treatment by weeks for B) and Bonferroni post hoc test for multiple comparisons; ^#^*p* < 0.05, ^##^*p* < 0.01, and ^###^*p* < 0.001 for 5xFAD-IgG2b vs. 5xFAD-5B8.

